# Characterization of a maintainable human myeloid model of VEXAS syndrome with enhanced TNF-induced cell death and DAMP release

**DOI:** 10.64898/2026.09.07.749759

**Authors:** Yuma Sakamoto, Nobuyasu Baba, Masanori Iseki, Eisei Kondo, Makoto Matsuyama, Tomoe Kobayashi, Yurika Shimizu, Shuji Mori, Tatsuo Ito, Tomoyuki Mukai

## Abstract

**Background:** VEXAS syndrome is an adult-onset autoinflammatory disorder caused by somatic *UBA1* mutations. *UBA1*-mutated myeloid cells have been reported to exhibit increased susceptibility to inflammatory cell death; however, the mechanisms underlying this phenotype and the extracellular consequences of enhanced cell death remain incompletely understood.

**Methods:** Disease-associated *UBA1* M41 mutations were introduced into U937 cells by CRISPR/Cas9-mediated genome editing. The resulting cells were characterized by genomic, molecular, and functional analyses, with particular focus on *UBA1*-associated cellular phenotypes, cell death signaling, and extracellular release of damage-associated molecular patterns (DAMPs).

**Results:** Genome editing yielded *UBA1*-mutated cells that could be maintained under standard culture conditions while recapitulating multiple molecular and cellular features associated with VEXAS syndrome. Long-read sequencing revealed the intended *UBA1* M41 mutation on one allele and a CRISPR/Cas9-induced on-target genomic deletion on the other. The mutant cells exhibited reduced UBA1b and increased UBA1c expression, prominent cytoplasmic vacuolization, reduced proliferative capacity, and increased basal cell death. They also showed enhanced susceptibility to TNF-induced cell death under conditions favoring either apoptosis or necroptosis. Mechanistic analyses demonstrated enhanced apoptotic and necroptotic signaling, together with increased basal abundance of RIPK1, RIPK3, and MLKL. Pharmacological inhibition of RIPK1, RIPK3, or MLKL attenuated membrane permeabilization under caspase-inhibited conditions. Enhanced cell death was accompanied by increased extracellular release of ATP, HMGB1, and S100A8/A9.

**Conclusions:** We characterized a maintainable *UBA1*-mutated human myeloid model that recapitulates multiple molecular and cellular features associated with VEXAS syndrome. Using this model, we showed that UBA1 dysfunction is associated with heightened susceptibility to TNF-dependent apoptotic and necroptotic cell death and increased extracellular release of multiple DAMPs. These findings provide insight into the cellular consequences of UBA1 dysfunction and establish a tractable experimental platform for further mechanistic studies of VEXAS syndrome.

**Key points:**

- We characterized a maintainable *UBA1*-mutated human myeloid model that recapitulates multiple molecular and cellular features associated with VEXAS syndrome.
- *UBA1*-mutated cells showed heightened susceptibility to TNF-dependent apoptotic and necroptotic cell death, accompanied by increased extracellular DAMP release.

## Introduction

VEXAS (vacuoles, E1 enzyme, X-linked, autoinflammatory, somatic) syndrome is a severe adult-onset autoinflammatory disease caused by acquired somatic mutations in *UBA1*, which encodes the ubiquitin-activating enzyme E1 [1]. This disease is characterized by recurrent fever and multisystem inflammation and is associated with poor clinical outcomes and limited responses to currently available therapies [2–7]. Despite increasing recognition of its clinical features, the molecular mechanisms driving inflammation in VEXAS syndrome remain incompletely understood. Recent studies have shown that *UBA1*-mutated myeloid cells exhibit altered inflammatory signaling and dysregulated cytokine responses [8, 9]. In addition, *UBA1*-mutated macrophages and monocytes have been reported to display increased susceptibility to TNF-induced cell death [10, 11]. However, the signaling mechanisms and the extracellular consequences of enhanced cell death remain incompletely understood.

At the molecular level, UBA1 is expressed as the nuclear UBA1a isoform and the cytoplasmic UBA1b isoform. The AUG codon encoding methionine 41 (M41) serves as the translation initiation site for the UBA1b isoform. Pathogenic M41 mutations impair translation initiation at this site, reducing UBA1b expression and increasing expression of UBA1c, a catalytically impaired isoform initiated from the downstream AUG codon encoding methionine 67 (M67) [1]. UBA1 mutations have also been associated with impaired ubiquitination and activation of proteostatic stress pathways, while recent studies have implicated RIPK1-dependent signaling in the enhanced TNF-induced cell-death phenotype [2, 4, 10–12]. Several genome-edited models of VEXAS syndrome have recently been developed, including hematopoietic stem/progenitor and myeloid cell systems [10–14]. However, maintainable human myeloid models that permit repeated biochemical and functional analyses of cell-death signaling remain limited.

Here, we introduced disease-associated *UBA1* M41 mutations into U937 cells to investigate the cellular consequences of UBA1 dysfunction. This approach yielded maintainable *UBA1*-mutated cells that recapitulated several VEXAS-associated features. Using these cells, we characterized TNF-induced apoptotic and necroptotic signaling, real-time cell-death responses, and the extracellular release of damage-associated molecular patterns (DAMPs) accompanying enhanced cell death.

## Materials and Methods

### Reagents and antibodies

The reagents and antibodies used in this study, including stimulants, inhibitors, neutralizing agents, and antibodies for immunoblotting, are listed in Supplementary Table S1.

### Generation of *UBA1* M41-mutated cells

Human male-derived monocytic U937 cells (CRL-1593.2; ATCC, Manassas, VA, USA) were used for genome-editing experiments targeting disease-associated *UBA1* M41 variants. In the initial series, parental U937 cells were subjected to CRISPR/Cas9-mediated genome editing followed by single-cell sorting to generate *UBA1*-mutated clones. In parallel, WT control clones were generated by the same electroporation and single-cell sorting procedures but without genome-editing reagents. Together, the *UBA1*-mutated and WT control clones constituted the initial parental-population-derived series (hereafter referred to as the bulk-derived series). To minimize the influence of pre-existing genetic heterogeneity, one WT control clone (WT #7) was selected as a common parental clone for a second round of genome editing to generate an independent single-clone-derived mutant series. The generated mutants were as follows: M41V (p.Met41Val, c.121A>G), M41T (p.Met41Thr, c.122T>C), and M41L (p.Met41Leu, c.121A>C). The mutations were introduced into exon 3 of the *UBA1* coding sequence for M41V, M41T, and M41L. Synthetic single-guide RNA (sgRNA), recombinant GFP-labeled Cas9 protein, and single-stranded donor DNAs were obtained from Integrated DNA Technologies Inc. (Coralville, IA, USA). Each donor DNA consisted of an 82-nt single-stranded oligonucleotide centered on the intended M41 nucleotide substitution and was used as a template for homology-directed repair (HDR). Electroporation of the ribonucleoprotein complexes, donor DNAs, and the HDR enhancer was performed using a NEPA21 electroporator (Nepa Gene, Ichikawa, Chiba, Japan). The sgRNA targeting sequence was designed as follows: 5′-GCCGTTCTTGGCCATTCCCT-3′.

Two days after electroporation, GFP-positive live single cells from the genome-editing conditions were sorted into 96-well plates using a FACSAria III (BD Biosciences, San Jose, CA, USA) to establish clonal cell lines. In parallel, live single cells from the WT control conditions were sorted using the same instrument without GFP-based selection. Once the single-cell-derived clones had sufficiently expanded, genomic DNA was extracted. The presence of the targeted mutations was confirmed by PCR amplification of a 408-bp genomic region encompassing the *UBA1* M41 site, followed by Sanger sequencing. Off-target candidate sequences were also checked using COSMID (CRISPR search with mismatches, insertions, and/or deletions; https://crispr.bme.gatech.edu/). The top three predicted off-target sites were examined by PCR and Sanger sequencing. The single-clone-derived mutant series was used for the principal analyses, whereas the independently established bulk-derived series was used for validation analyses. The identity of the parental U937 cells was confirmed by short tandem repeat (STR) profiling before genome-editing experiments. Cells were routinely tested and confirmed negative for mycoplasma contamination.

### DNA sequencing

PCR products were purified using ExoSAP-IT™ (Thermo Fisher Scientific, Waltham, MA, USA) to remove residual primers and nucleotides and subjected to Sanger sequencing by a commercial service (FASMAC Co., Ltd., Atsugi, Kanagawa, Japan) to confirm the presence of the intended mutations. The primer sequences are listed in Supplementary Table S2.

### Cell culture

U937 cells were maintained in RPMI 1640 medium supplemented with 10% fetal bovine serum (FBS), 100 U/mL penicillin, and 100 µg/mL streptomycin (Thermo Fisher Scientific) at 37°C in a humidified atmosphere containing 5% CO□. Cells were treated with TNF-α, LCL161, z-VAD-fmk, or other reagents under the indicated experimental conditions.

### Nanopore long-read sequencing of the *UBA1* genomic locus and transcripts

Total RNA samples isolated from U937 cells were reverse-transcribed into cDNA using the PrimeScript RT Reagent Kit (Takara Bio) with random primers. For genomic DNA analysis, long-range PCR was performed using genomic DNA extracted from U937 cells. The primer sequences are listed in Supplementary Table S2. The resulting cDNA and genomic PCR amplicons were quantified using the QuantiFluor dsDNA System (Promega, Madison, WI, USA). Sequencing libraries were prepared using the Rapid Barcoding Kit V14 (SQK-RBK114; Oxford Nanopore Technologies), followed by purification with Agencourt AMPure XP beads (Beckman Coulter, Brea, CA, USA). The libraries were loaded onto PromethION R10.4.1 flow cells (FLO-PRO114M; Oxford Nanopore Technologies) and sequenced using a PromethION 2 Solo device. Basecalling was performed using Dorado (v7.9.8) in super-accuracy mode, and reads with quality scores <9 were excluded. Processed reads were aligned to the human reference genome (GRCh38) using minimap2 (v2.2). The resulting BAM files were sorted and indexed with samtools (v1.11). Visualization and manual inspection of aligned reads and transcript structures were performed using Integrative Genomics Viewer (IGV, v2.9.4).

### Quantitative PCR analysis

Total RNA was extracted from cultured cells using RNAiso Plus (Takara Bio), and cDNA was synthesized using the PrimeScript RT Reagent Kit (RR037, Takara Bio). Quantitative PCR (qPCR) was performed using TB Green PCR Master Mix (RR820, Takara Bio) on a QuantStudio 5 Real-Time PCR System (Thermo Fisher Scientific), as described previously [15]. The expression levels of *UBA1*, *RIPK1*, *RIPK3*, *MLKL*, and *CASP3* were analyzed. For UBA1, three primer pairs targeting the exon 1– 2 junction/exon 2, exons 5–6, and the exon 15–16 junction/exon 16 were used to assess transcript levels at different regions of the gene. Relative mRNA expression levels were normalized to *HPRT1* using the 2^-ΔΔCt^ method and expressed relative to the indicated control samples. *GAPDH* was also evaluated as an internal control and yielded comparable results. Primer sequences are provided in Supplementary Table S2. All reactions generated single-peak dissociation curves, confirming amplification specificity.

### Morphological analysis by May-Grünwald-Giemsa staining

Cells were suspended at 1 × 10^5^ cells/200 μL in culture medium and cytocentrifuged onto glass slides using a Cytospin 3 cytocentrifuge (Thermo Fisher Scientific) at 450 rpm for 5 min. The slides were immediately air-dried and subjected to May-Grünwald-Giemsa staining. Briefly, slides were stained with May-Grünwald solution (Muto Pure Chemicals Co., Ltd., Tokyo, Japan) for 5 min, followed by Giemsa solution (Muto Pure Chemicals Co., Ltd.) diluted in phosphate buffer for 20 min. After rinsing with distilled water and air-drying, stained cells were observed and photographed under a light microscope (IX83, Olympus, Tokyo, Japan).

### Cell proliferation assay

Cell proliferation was evaluated using a Cell Counting Kit-8 (CCK-8) (Dojindo Laboratories, Kumamoto, Japan). Cells were seeded into 96-well plates at 5 × 10^3^ cells/well. At 24, 48, and 72 h after seeding, 10 µL of CCK-8 solution was added to each well and incubated for 3 h at 37°C. Absorbance at 450 nm was measured using a Varioskan microplate reader (Thermo Fisher Scientific).

### Lactate dehydrogenase (LDH) release assay

Cell death was assessed by measuring LDH release into culture supernatants using the Cytotoxicity LDH Assay Kit-WST (Dojindo Laboratories) according to the manufacturer’s instructions. Cells were seeded at 2 × 10^4^ cells/well and cultured in RPMI 1640 medium supplemented with 1% FBS and treated with the indicated reagents. Maximum LDH release was determined after complete cell lysis using the provided lysis buffer. LDH release was expressed as a percentage of the maximum release, and absorbance was measured at 490 nm using a Varioskan microplate reader (Thermo Fisher Scientific). ΔLDH release was calculated by subtracting the value of the corresponding untreated control.

### Flow cytometry

Cell death was assessed by flow cytometry using Annexin V-APC and propidium iodide (PI) co-staining. Cells were washed twice with ice-cold PBS and resuspended in 1× Annexin V binding buffer (BD Biosciences) at 1 × 10^6^ cells/mL. Aliquots of 100 μL (1 × 10^5^ cells) were stained with 5 µL of Annexin V-APC and 5 µL of PI for 15 min at 25°C in the dark. After incubation, 400 µL of 1× Annexin V binding buffer was added to each sample, and the samples were analyzed using a CytoFLEX flow cytometer (Beckman Coulter). Data were analyzed with the aid of FlowJo software (version 10, BD Biosciences). Annexin V/PI staining patterns were quantified.

### Cell cycle

Cell cycle distribution was analyzed by flow cytometry based on DNA content. Cells were harvested, washed with PBS, and resuspended in 250 µL staining solution containing PI (100 µg/mL), RNase A (800 µg/mL), Triton X-100 (0.1%), and sodium azide (0.1 mg/mL) in PBS. After incubation for 30 min at 37°C in the dark, DNA content was analyzed using a CytoFLEX flow cytometer (Beckman Coulter). The sub-G1 fraction and the G0/G1, S, and G2/M phases were quantified with the aid of FlowJo software (version 10, BD Biosciences) after exclusion of doublets.

### RealTime-Glo^TM^ MT Cell Viability Assay and CellTox Green Cytotoxicity Assay

Cell viability and cell death were assessed using a combination of the RealTime-Glo™ MT Cell Viability Assay and the CellTox Green Cytotoxicity Assay (Promega) according to the manufacturer’s instructions. WT and *UBA1*-mutated U937 cells were seeded into white 96-well plates at a density of 5 × 10^3^ cells/well and allowed to rest for 2 h at 37°C. RealTime-Glo™ MT substrates and CellTox Green reagent were then added to each well, followed by treatment with the indicated reagents. Luminescence, reflecting the metabolic activity of viable cells, and fluorescence, reflecting loss of plasma membrane integrity, were measured at the indicated time points over a 72 h incubation period using a Varioskan microplate reader (Thermo Fisher Scientific).

### RealTime-Glo™ Annexin V Apoptosis and Necrosis Assay

Apoptosis and necrosis were monitored using the RealTime-Glo™ Annexin V Apoptosis and Necrosis Assay (Promega) according to the manufacturer’s instructions. WT and *UBA1*-mutated U937 cells were seeded into white 96-well plates at the indicated density. Assay reagents, including Annexin V NanoBiT® substrate and the necrosis detection dye, were added to each well, and cells were treated with the indicated reagents under the specified conditions. Luminescence (Annexin V signal, reflecting phosphatidylserine exposure) and fluorescence (membrane-impermeant DNA dye signal, reflecting loss of membrane integrity) were measured at regular intervals using a Varioskan microplate reader (Thermo Fisher Scientific) maintained at 37°C with 5% CO.

### RealTime-Glo™ Extracellular ATP Assay

Extracellular ATP levels were monitored in culture wells using the RealTime-Glo™ Extracellular ATP Assay (Promega) according to the manufacturer’s instructions. WT and *UBA1*-mutated cells were seeded in white 96-well plates at 2 × 10^4^ cells/well. After a 6 h culture, 4× assay reagent (prepared according to the manufacturer’s instructions by reconstituting with the cell culture medium) was directly added to each well at a final concentration of 25% (v/v), followed by treatment with the indicated reagents. Luminescence, reflecting ATP levels, was measured at regular intervals using a Varioskan microplate reader (Thermo Fisher Scientific) maintained at 37°C with 5% CO.

### Western blotting

Western blotting was performed using whole-cell lysates prepared from cultured cells, as described previously [16]. Cells were washed with ice-cold PBS and lysed in RIPA buffer (R0278; Merck Millipore, Burlington, MA, USA) supplemented with protease and phosphatase inhibitor cocktails (P8340, P5726, and P0044; Merck Millipore), added immediately before use. Protein concentrations were determined using a BCA Protein Assay Kit (23227, Thermo Fisher Scientific).

Proteins were resolved by sodium dodecyl sulfate-polyacrylamide gel electrophoresis (SDS-PAGE) and transferred onto PVDF membranes. Membranes were blocked with 5% skim milk in Tris-buffered saline containing 0.1% Tween-20 (TBS-T) for non-phosphorylated proteins or with 5% bovine serum albumin (BSA) in TBS-T for phosphorylated proteins. After blocking, membranes were incubated with the indicated primary antibodies, followed by the appropriate horseradish peroxidase (HRP)-conjugated secondary antibodies. Protein bands were detected using Clarity Max Western ECL chemiluminescent substrate (Bio-Rad, Hercules, CA, USA) and visualized using an Amersham Imager 680 (GE Healthcare, Little Chalfont, UK). β-actin was used as a loading control.

### Generation of a monoclonal antibody against UBA1a/b

A peptide corresponding to amino acids 42-60 (AKNGSEADIDEGLYSRQLY) of human UBA1a/b was synthesized and conjugated to keyhole limpet hemocyanin (KLH) (Sigma-Aldrich Japan, Tokyo, Japan). A rat monoclonal antibody against UBA1a/b was generated using the iliac lymph node method as previously described [17]. Rats were immunized with the peptide-KLH conjugate, and lymphocytes isolated from the iliac lymph nodes were fused with myeloma cells to generate hybridomas. Culture supernatants were screened by a solid-phase ELISA and western blotting, and hybridomas specifically recognizing UBA1a/b but not UBA1c were selected and subcloned by limiting dilution. Culture supernatants from the established hybridomas were used for western blotting.

### Enzyme-Linked Immunosorbent Assay (ELISA)

Concentrations of TNF, IL-1β, IL-6, IL-8, and S100A8/A9 in culture supernatants were measured using DuoSet ELISA Development Systems (R&D Systems, Minneapolis, MN, USA), whereas HMGB1 was measured using an HMGB1 ELISA kit (Shino-Test Corporation, Tokyo, Japan). Culture supernatants were centrifuged to remove cellular debris. All assays were performed according to the manufacturers’ instructions, and analyte concentrations were calculated from the corresponding standard curves.

### NF-κB luciferase reporter assays

U937 NFκB-LUC2 cells (CRL-1593.2-NFkB-LUC2; ATCC, Manassas, VA, USA), which stably express a luciferase reporter under the control of an NF-κB-responsive promoter, were used as reporter cells. For conditioned-medium experiments, culture supernatants were collected from WT or *UBA1*-mutated U937 cells after 24 h under the indicated conditions, centrifuged to remove cells, and 50 μL was added to reporter cells seeded at 7.5 × 10^3^ cells/well. Where indicated, conditioned media were pre-incubated with etanercept (10 μg/mL) before addition to reporter cells. For direct co-culture experiments, reporter cells were co-cultured with WT or *UBA1*-mutated U937 cells at 3.5 × 10^3^ cells of each population per well. NF-κB reporter activity was measured after the indicated incubation periods using the ONE-Glo Luciferase Assay System (Promega), and luminescence was quantified with a Varioskan microplate reader (Thermo Fisher Scientific).

### Data presentation

Data are presented as mean ± SD where applicable. The numbers of independent experiments and technical replicates are specified in the figure legends. Because only a limited number of independently derived genome-edited clones were available, inferential statistical testing at the genotype level was not performed. Graphs were generated using GraphPad Prism version 9 (GraphPad Software, Boston, MA, USA).

## Results

### 1. Generation and initial characterization of *UBA1*-mutated U937 cells

To investigate the cellular consequences of disease-associated *UBA1* mutations, we introduced *UBA1* M41 mutations into U937 cells using CRISPR/Cas9-mediated genome editing. The CRISPR/Cas9 editing design, including the target sequences and corresponding gRNA design, is illustrated in Figure 1A. In the initial series, *UBA1*-mutated clones were generated by CRISPR/Cas9-mediated genome editing of parental U937 cells followed by single-cell sorting, whereas WT control clones were established in parallel by the same electroporation and single-cell sorting procedures without genome-editing reagents (Figure 1B). Together, these clones constituted the parental-population-derived series (bulk-derived series). Because independently isolated clones from the parental population could differ in their pre-existing genetic backgrounds, we considered that such heterogeneity might complicate precise genotype–phenotype comparisons. We therefore selected one WT control clone (WT #7) from this initial series and used it as a common parental clone for a second round of genome editing. This generated an independent single-clone-derived mutant series in which the mutant clones and WT control shared a common clonal background (Figure 1B). The single-clone-derived series was consequently used for the principal mechanistic analyses, whereas the independently generated bulk-derived series was used to assess the reproducibility of the observed phenotypes.

**Figure 1.**
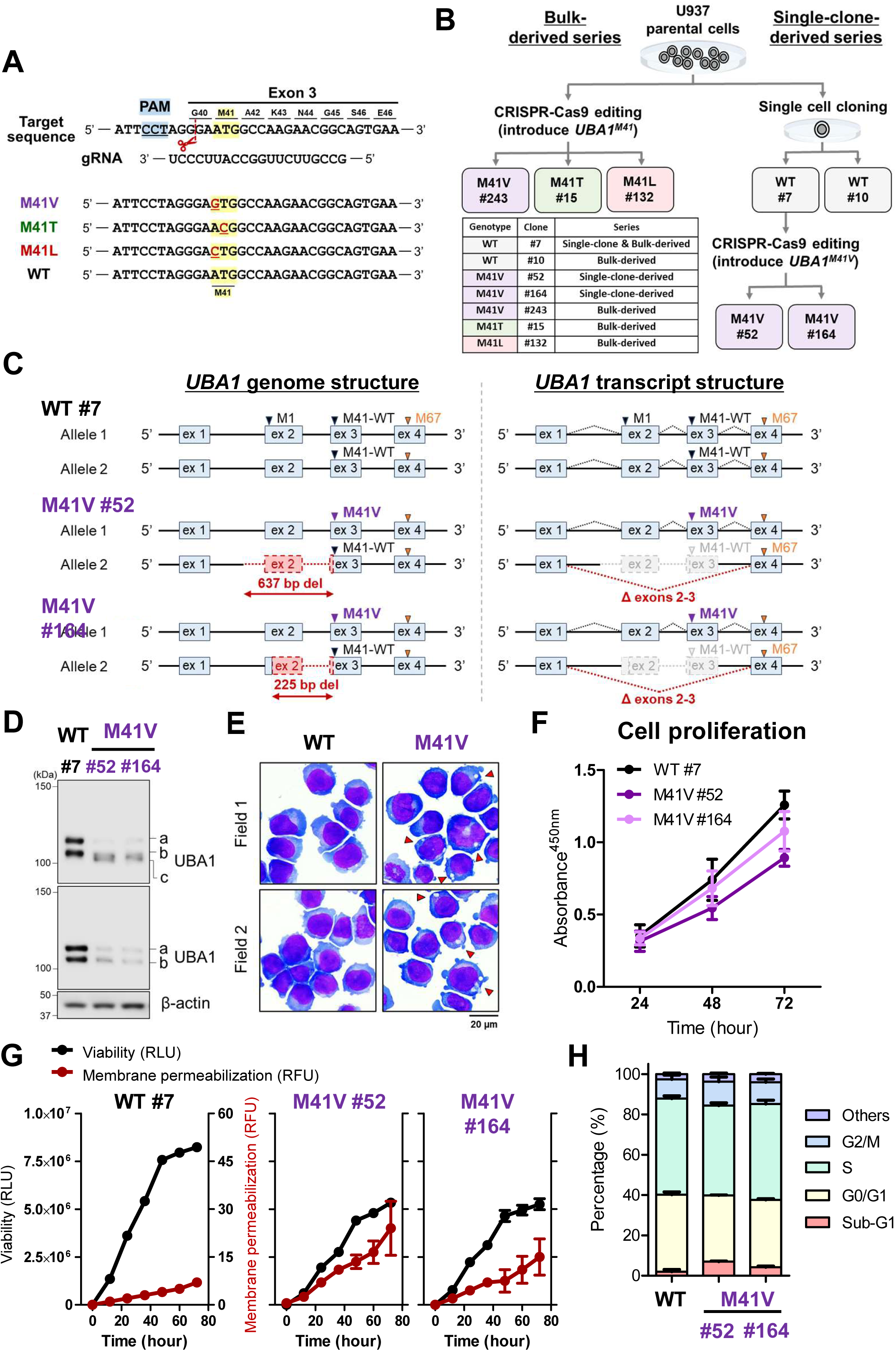
Genomic and cellular characterization of *UBA1*-mutated U937 cells. **(A)** Schematic representation of CRISPR/Cas9-mediated genome editing targeting *UBA1* exon 3. The target sequence, protospacer-adjacent motif (PAM), guide RNA sequence, and nucleotide substitutions introduced to generate the M41V, M41T, and M41L variants are shown. **(B)** Workflow for establishing bulk-derived and single-clone-derived *UBA1*-mutated U937 cell series. In the initial bulk-derived series, *UBA1*-mutated clones were generated by genome editing followed by single-cell sorting, whereas WT control clones were generated in parallel using the matched procedure without genome-editing reagents. Subsequently, single-clone-derived M41V clones (#52 and #164) were generated from the parental WT clone (#7). The clones used in the present study are summarized. **(C)** Genomic and transcript structures of WT #7 and M41V clones. Genome structures were determined by long-range PCR followed by nanopore long-read sequencing. Both M41V clones carried the intended M41V substitution in one allele and a 637-bp (#52) or 225-bp (#164) deletion encompassing exon 2 in the other allele. Transcript analysis identified transcripts from the deletion allele lacking exons 2 and 3, with exon 1 joined directly to exon 4. **(D)** Immunoblot analysis of UBA1 isoforms. The upper blot was probed with an antibody recognizing UBA1a, UBA1b, and UBA1c, whereas the lower blot was probed with an antibody recognizing UBA1a and UBA1b. β-actin served as a loading control. **(E)** Representative May-Grünwald-Giemsa-stained images of WT and M41V cells. Arrowheads indicate cytoplasmic vacuoles. Scale bar, 20 μm. **(F)** Cell proliferation was evaluated using the Cell Counting Kit-8 assay. Data are presented as mean ± SD of four independent experiments, each performed in triplicate. **(G)** Cell viability and plasma membrane permeabilization were monitored for 72 h using the RealTime-Glo ^MT^ Cell Viability Assay and CellTox Green Cytotoxicity Assay, respectively. Data are presented as mean ± SD of duplicate wells of a representative experiment. **(H)** Cell-cycle distribution was analyzed by flow cytometry. Data are presented as mean ± SD of two independent experiments. RLU: Relative Light Units, RFU: Relative Fluorescence Units.

Extensive screening was required to obtain viable *UBA1*-mutated clones suitable for further analysis. Eleven independent genome-editing experiments were performed, and GFP-positive single cells were individually sorted by flow cytometry into a total of 65,184 wells of 96-well plates. Among 3,456 genotyped clones, 72 were initially identified as carrying the intended M41 substitution at the target locus, and 26 showed no detectable sequence alterations at the predicted off-target sites. A subset of these clones was selected for further analyses after additional genetic and cellular quality assessment. Clones showing extensive genomic abnormalities on initial screening, unexpected insertions, or atypical UBA1 isoform expression were excluded. Accordingly, five independently derived mutant clones were selected for further analyses (Supplementary Table S3). These selected clones were subsequently subjected to detailed genomic characterization.

The introduced *UBA1* mutations were initially confirmed by Sanger sequencing. Although U937 cells were originally established from a male patient, this cell line carries two X chromosomes and therefore harbors two *UBA1* alleles [18]. To comprehensively characterize the structure of both edited alleles, we designed multiple genomic and transcript PCR assays spanning the *UBA1* locus and performed conventional PCR and RT-PCR (Supplementary Figures S1A, S1B, and S2A), followed by nanopore long-read sequencing (Supplementary Figures S1C, S2C, S3, and S4). Long-read sequencing revealed that CRISPR/Cas9-mediated cleavage at the target site was accompanied by genomic deletions extending toward the 5′ region of the *UBA1* locus, with the selected surviving clones carrying the intended M41 mutation on one allele and a genomic deletion on the other (Figure 1C and Supplementary Figures S2B, S3, and S4). At the transcript level, long-read sequencing identified transcripts derived from the deletion allele that lacked exons 2 and 3, with exon 1 joined directly to exon 4. Consistent with this structure, read coverage across exons 2 and 3 was reduced to approximately half that observed in WT cells, whereas coverage of exons 1, 4, and 5 was largely preserved (Supplementary Figures S1C and S2C). Quantitative RT-PCR using primers targeting the exon 1–2 junction and exon 2 further demonstrated reduced expression of transcripts containing this region, whereas no apparent differences were observed using primers targeting exons 5–6 or the exon 15–16 junction and exon 16 (Supplementary Figure S1D).

We next examined whether the surviving genome-edited *UBA1*-mutated cells recapitulated molecular and cellular features associated with VEXAS syndrome. *UBA1*-mutated cells exhibited reduced UBA1b and increased UBA1c expression, consistent with the characteristic UBA1 isoform alterations observed in VEXAS syndrome (Figure 1D and Supplementary Figure S2D). May-Grünwald-Giemsa staining further demonstrated prominent cytoplasmic vacuolization in the mutated cells (Figure 1E). The *UBA1*-mutated cells showed reduced proliferative capacity and evidence of increased basal membrane permeabilization (Figure 1F, G and Supplementary Figure S2E, F). Despite increased basal cell death, the *UBA1*-mutated cells remained maintainable under standard culture conditions. Cell-cycle analysis additionally demonstrated an increased sub-G1 fraction and a reduced S-phase fraction in *UBA1*-mutated cells (Figure 1H and Supplementary Figure S2G).

Collectively, these findings indicate that the *UBA1*-mutated U937 cells remained maintainable while recapitulating several molecular and cellular features associated with VEXAS syndrome, providing a tractable system for subsequent functional analyses.

### 2. *UBA1*-mutated cells exhibit enhanced basal and TNF-induced cell death

The initial characterization suggested impaired cell growth and survival in *UBA1*-mutated cells. We therefore first examined spontaneous cell death under basal culture conditions. *UBA1*-mutated cells showed a modest increase in basal cell death, as indicated by increased extracellular LDH release and a higher proportion of Annexin V-positive cells compared with WT cells (Figure 2A, B and Supplementary Figure S5A, B).

**Figure 2.**
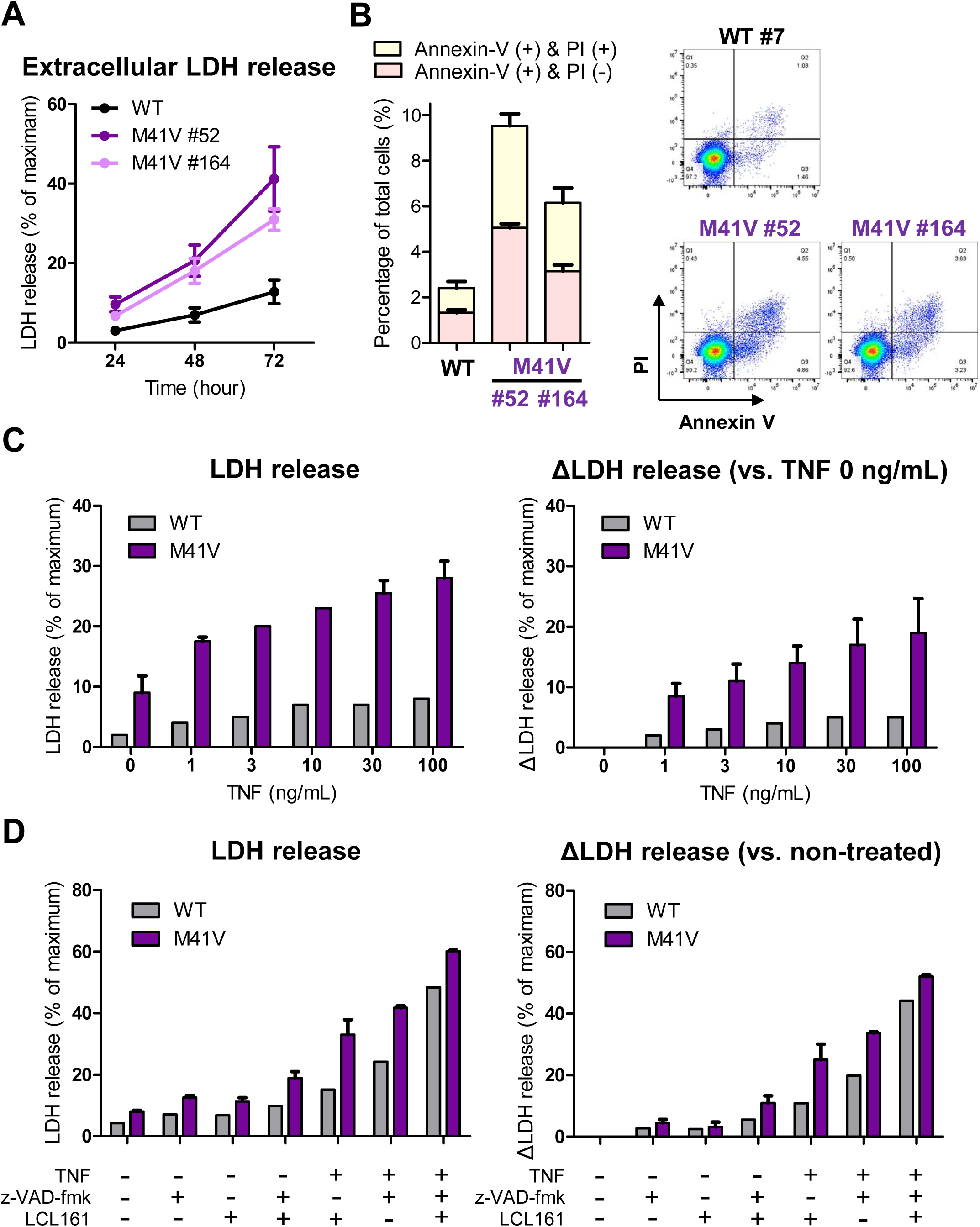
*UBA1*-mutated cells exhibit increased sensitivity to TNF-induced cell death. **(A)** Spontaneous cell death under basal culture conditions was evaluated by measuring extracellular lactate dehydrogenase (LDH) release over 72 h. Data are presented as mean ± SD of four independent experiments, each performed in triplicate. **(B)** Cell death under basal culture conditions was analyzed after 24 h by Annexin V/propidium iodide (PI) staining and flow cytometry. Representative flow cytometry plots are shown on the right. Bar graphs represent the percentages of Annexin V-positive/PI-negative and Annexin V-positive/PI-positive cells. Data are presented as mean ± SD of three independent experiments. **(C)** Cells were stimulated with the indicated concentrations of TNF for 72 h, and extracellular LDH release was measured. The left panel shows LDH release expressed as a percentage of maximum LDH release, and the right panel shows ΔLDH release relative to cells treated with 0 ng/mL TNF. Data are presented as mean ± SD of two independent experiments, each performed in triplicate. **(D)** Cells were treated with TNF (10 ng/mL), z-VAD-fmk (10 μM), LCL161 (250 nM), or the indicated combinations for 72 h, followed by measurement of extracellular LDH release. The left panel shows LDH release expressed as a percentage of maximum LDH release, and the right panel shows ΔLDH release relative to non-treated cells. Data are presented as mean ± SD of duplicate wells of a representative experiment.

Because increased susceptibility to TNF-induced cell death has been reported in VEXAS syndrome [10, 11], we next examined the response of *UBA1*-mutated cells to TNF. Across a range of TNF concentrations, *UBA1*-mutated cells showed greater LDH release than WT cells, including at low TNF concentrations (Figure 2C). Similar increases in basal cell death and TNF sensitivity were observed in the independently generated bulk-derived series (Supplementary Figure S5A-C).

We next examined how caspase inhibition or cIAP antagonism affected cell death in *UBA1*-mutated cells, both in the absence and presence of TNF. Cells were treated with the pan-caspase inhibitor z-VAD-fmk or the cIAP antagonist LCL161, alone or in combination with TNF (Figure 2D). In the absence of exogenous TNF, only small genotype-associated differences in LDH release were observed under conditions of caspase inhibition or cIAP antagonism. These differences became more evident in the presence of TNF, with prominent genotype-associated differences observed under both TNF plus z-VAD-fmk and TNF plus LCL161 conditions (Figure 2D).

Taken together, these findings indicate that *UBA1*-mutated cells exhibit a modest increase in basal cell death and heightened sensitivity to TNF-induced cell death across multiple signaling contexts.

### 3. *UBA1*-mutated cells exhibit enhanced apoptotic and necroptotic signaling downstream of TNF

To determine the signaling basis of the heightened cell-death susceptibility of *UBA1*-mutated cells, we examined the engagement of apoptotic and necroptotic pathways under the conditions identified in Figure 2. We first asked whether the enhanced cell death observed under caspase inhibition was accompanied by increased activation of the RIPK1–RIPK3–MLKL necroptotic pathway. Following TNF plus z-VAD-fmk treatment, phosphorylation of RIPK1, RIPK3, and MLKL was more evident in *UBA1*-mutated cells than in WT cells (Figure 3A). In addition, total RIPK1, RIPK3, and MLKL protein levels were higher in *UBA1*-mutated cells even before stimulation (Figure 3A). Thus, under caspase-inhibited conditions, the heightened cell-death phenotype of *UBA1*-mutated cells was accompanied by enhanced signaling through core components of the necroptotic pathway, together with increased basal abundance of these proteins.

**Figure 3.**
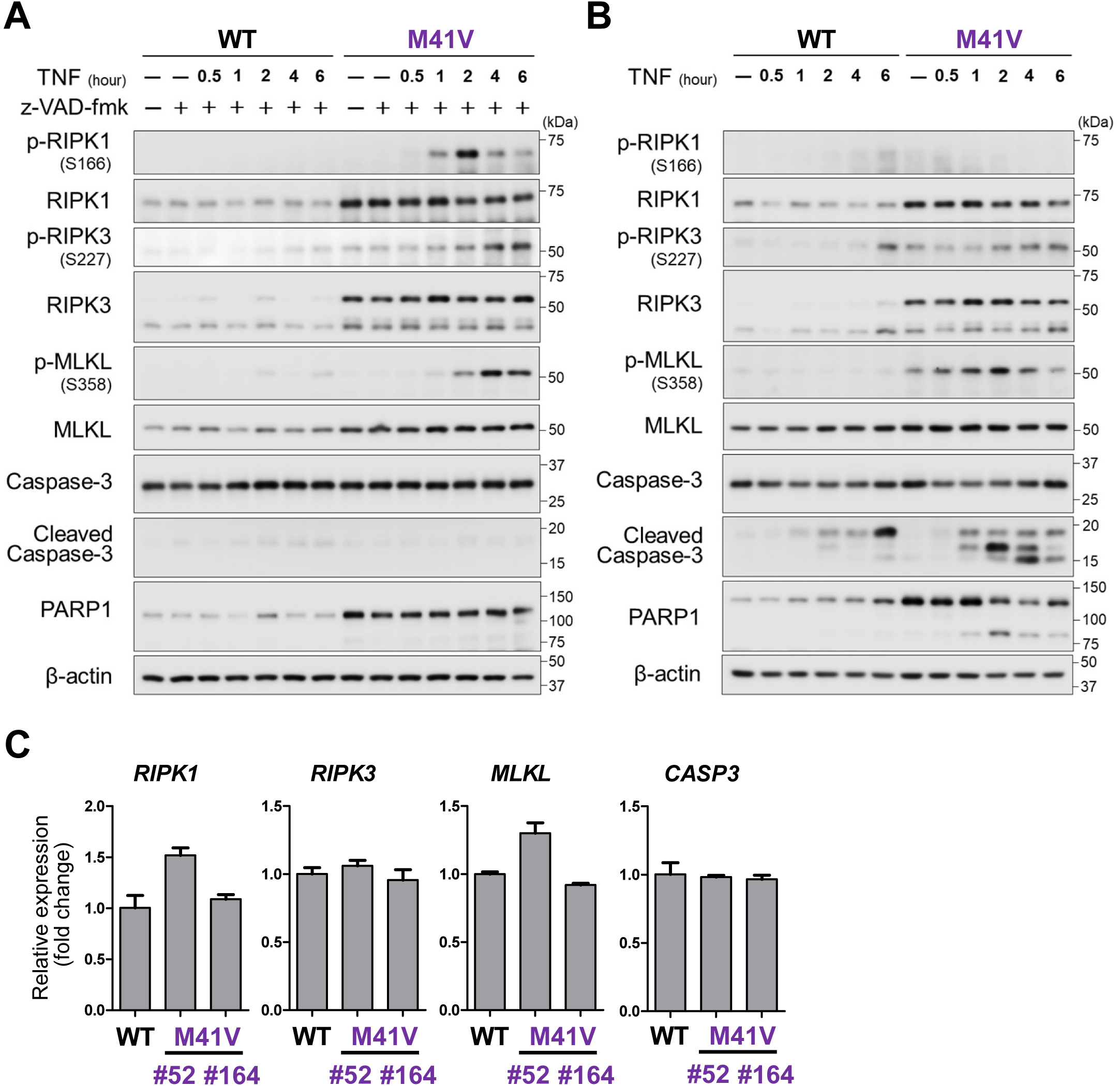
Enhanced cell death signaling in *UBA1*-mutated cells following TNF stimulation. **(A)** Cells were stimulated with TNF (10 ng/mL) in the presence of z-VAD-fmk (10 μM) for the indicated times. Cell lysates were analyzed by immunoblotting for phosphorylated and total RIPK1, RIPK3, and MLKL, as well as caspase-3, cleaved caspase-3, and PARP1. β-actin served as a loading control. **(B)** Cells were stimulated with TNF (10 ng/mL) for the indicated times. Cell lysates were analyzed by immunoblotting for phosphorylated and total RIPK1, RIPK3, and MLKL, as well as caspase-3, cleaved caspase-3, and PARP1. β-actin served as a loading control. Full-length and cleaved caspase-3 detected with the same antibody are shown separately in (A) and (B) because different exposure times were used to visualize the cleaved form. Representative immunoblots are shown in (A–B). **(C)** Relative mRNA expression levels of *RIPK1*, *RIPK3*, *MLKL*, and *CASP3* were measured by quantitative PCR under basal culture conditions. Expression levels were normalized to *HPRT1*, and relative expression was calculated with WT set to 1. Representative data from technical duplicates are shown.

We next examined cell-death signaling following TNF stimulation alone, without pharmacological modulation of caspases or cIAPs. Under this condition, caspase-3 activation and PARP1 cleavage were more apparent in *UBA1*-mutated cells than in WT cells (Figure 3B). Modest phosphorylation of RIPK3 and MLKL was also observed in the mutant cells, whereas RIPK1 phosphorylation remained minimal (Figure 3B). Thus, TNF stimulation alone was associated primarily with enhanced apoptotic signaling, accompanied by modest activation of necroptotic signaling components.

To determine whether enhanced apoptotic signaling was also observed under another TNF-dependent cell-death condition, we examined cells treated with TNF plus the cIAP antagonist LCL161. Under this condition, *UBA1*-mutated cells exhibited more prominent cleavage of caspase-3 and PARP1 than WT cells (Supplementary Figure S6A), indicating enhanced apoptotic signaling under cIAP antagonism.

In contrast, canonical NF-κB activation following TNF stimulation was broadly comparable between WT and *UBA1*-mutated cells (Supplementary Figure S6B), indicating that the enhanced cell-death response was not accompanied by a corresponding increase in canonical NF-κB signaling.

Because several core necroptotic proteins were already increased before stimulation, we further examined whether basal differences extended more broadly to components of TNFR1-associated signaling. *UBA1*-mutated cells showed higher basal abundance of several TNFR1 signaling-associated proteins, including A20, cIAP1, TRADD, and FADD (Supplementary Figure S6C). Thus, genotype-associated differences were not restricted to stimulus-induced phosphorylation or cleavage but also involved the basal abundance of multiple components of the TNFR1 signaling machinery.

We further examined basal mRNA expression of cell death-related genes by quantitative RT-PCR. No consistent genotype-associated increases in the mRNA expression of *RIPK1*, *RIPK3*, *MLKL*, or *CASP3* were observed between WT and *UBA1*-mutated cells (Figure 3C).

Collectively, these findings demonstrate context-dependent enhancement of both apoptotic and necroptotic signaling downstream of TNF, together with altered basal abundance of multiple TNFR1-associated signaling proteins. Necroptotic signaling was particularly prominent under caspase-inhibited conditions, whereas enhanced apoptotic signaling was evident following TNF stimulation alone and under cIAP antagonism.

### 4. Enhanced cell death in *UBA1*-mutated cells involves RIPK1–RIPK3–MLKL signaling under caspase-inhibited conditions

Having identified enhanced apoptotic and necroptotic signaling, we next examined whether these molecular differences were reflected in cell-death responses over time. Phosphatidylserine exposure and loss of membrane integrity were simultaneously monitored using the RealTime-Glo™ Annexin V Apoptosis and Necrosis Assay. Compared with WT cells, *UBA1*-mutated cells showed greater increases in both Annexin V and membrane-permeabilization signals under several conditions, including TNF stimulation and caspase inhibition, with the most pronounced genotype-associated difference observed under TNF plus z-VAD-fmk conditions (Figure 4A). Under TNF plus z-VAD-fmk treatment, both Annexin V and membrane-permeabilization signals increased substantially more in *UBA1*-mutated cells than in WT cells, indicating a greater extent of cell death under caspase-inhibited conditions. A similar tendency toward greater cell-death responses was observed under additional TNF-related conditions and in the independently generated bulk-derived series (Supplementary Figures S7 and S8).

**Figure 4.**
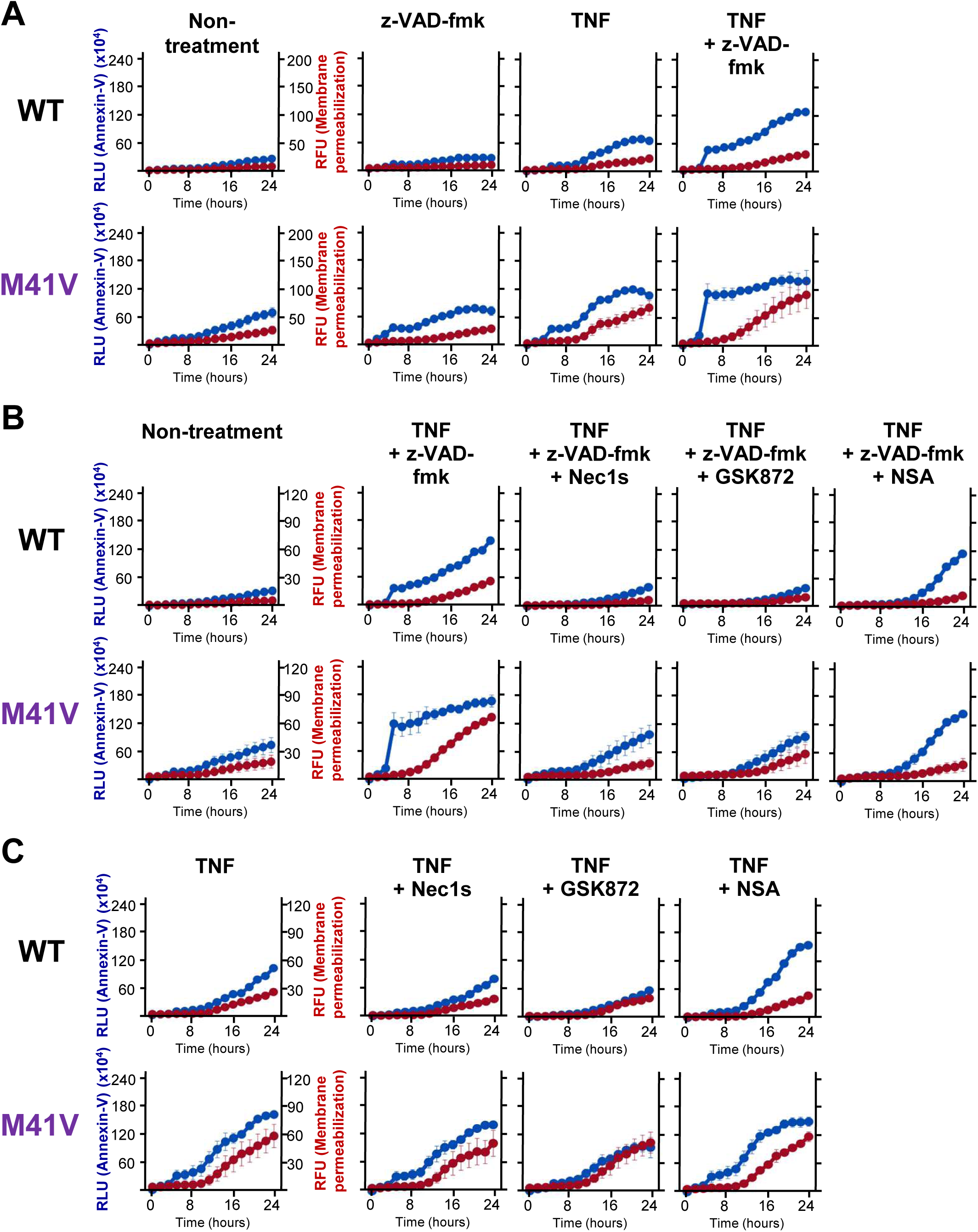
*UBA1*-mutated cells exhibit enhanced TNF-dependent cell death with RIPK1–RIPK3–MLKL pathway involvement under caspase inhibition. (A) Cells were stimulated with TNF (10 ng/mL), z-VAD-fmk (10 μM), or both, as indicated. Annexin V luminescence (blue) and membrane permeabilization-associated fluorescence (red), reflecting phosphatidylserine exposure and membrane permeabilization, respectively, were monitored for 24 h using the RealTime-Glo™ Annexin V Apoptosis and Necrosis Assay. **(B)** Cells were stimulated with TNF (10 ng/mL) plus z-VAD-fmk (10 μM) in the absence or presence of Nec-1s (10 μM), GSK872 (10 μM), or NSA (10 μM), as indicated. Annexin V luminescence (blue) and membrane permeabilization-associated fluorescence (red) were monitored for 24 h using the same assay. **(C)** Cells were stimulated with TNF (10 ng/mL) in the absence or presence of Nec-1s (10 μM), GSK872 (10 μM), or NSA (10 μM), as indicated. Annexin V luminescence (blue) and membrane permeabilization-associated fluorescence (red) were monitored for 24 h using the same assay. Representative data are shown in (A–C). RLU: Relative Light Units, RFU: Relative Fluorescence Units.

Because the biochemical analyses in Figure 3 showed enhanced RIPK1–RIPK3–MLKL signaling under TNF plus z-VAD-fmk conditions, we next asked whether this pathway functionally contributed to the increased loss of membrane integrity. Pharmacological inhibition of RIPK1, RIPK3, or MLKL with Nec-1s, GSK872, or NSA, respectively, substantially reduced the membrane-permeabilization signal induced by TNF plus z-VAD-fmk in *UBA1*-mutated cells (Figure 4B). The effects of these inhibitors on Annexin V signals were less pronounced than those on membrane permeabilization. These findings indicate that RIPK1–RIPK3–MLKL signaling contributes to the enhanced loss of membrane integrity under caspase-inhibited conditions.

We further examined whether the same pathway made a comparable contribution to cell death induced by TNF alone. In contrast, RIPK1, RIPK3, or MLKL inhibition had only limited effects on membrane permeabilization following TNF stimulation alone (Figure 4C), indicating a smaller contribution of the canonical necroptotic pathway under this condition.

Collectively, these real-time analyses demonstrate that *UBA1*-mutated cells exhibit greater cell-death responses under multiple TNF-related conditions. Under caspase-inhibited conditions, pharmacological inhibition of RIPK1, RIPK3, or MLKL markedly suppressed the enhanced membrane-permeabilization response, providing functional support for increased necroptotic cell-death execution in UBA1-mutated cells under caspase-inhibited conditions.

### 5. Enhanced cell death in *UBA1*-mutated cells is accompanied by increased extracellular DAMP release

To determine whether the enhanced cell-death phenotype of *UBA1*-mutated cells was accompanied by increased extracellular release of DAMPs, we first monitored extracellular ATP over time. Extracellular ATP levels were low under basal conditions but increased following z-VAD-fmk, TNF, or TNF plus z-VAD-fmk treatment, with the most prominent increase observed in *UBA1*-mutated cells under TNF plus z-VAD-fmk conditions (Figure 5A). A similar pattern was observed in the bulk-derived series (Supplementary Figure S9A).

**Figure 5.**
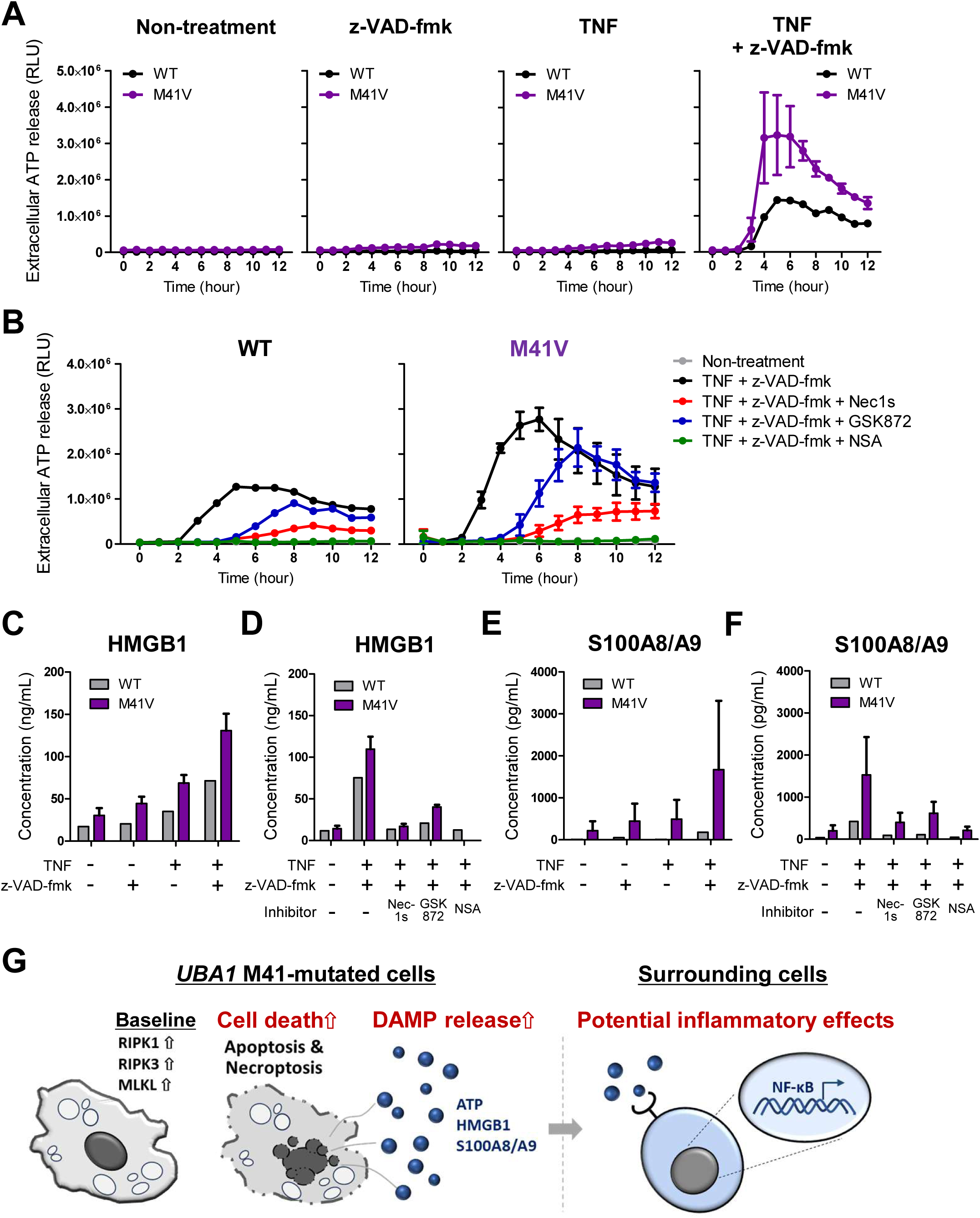
Enhanced cell death in *UBA1*-mutated cells is accompanied by increased extracellular DAMP release. **(A)** Cells were left untreated or stimulated with z-VAD-fmk (10 μM), TNF (10 ng/mL), or TNF (10 ng/mL) plus z-VAD-fmk (10 μM). Extracellular ATP levels were measured at the indicated time points over 12 h. Representative data from duplicate wells are shown. **(B)** Cells were stimulated with TNF (10 ng/mL) plus z-VAD-fmk (10 μM) in the presence or absence of Nec-1s (10 μM), GSK872 (10 μM), or NSA (10 μM). Extracellular ATP levels were measured at the indicated time points over 12 h. Representative data from duplicate wells are shown. **(C, E)** Cells were stimulated with z-VAD-fmk (10 μM), TNF (10 ng/mL), or TNF plus z-VAD-fmk (10 μM) for 24 h. Extracellular HMGB1 (C) and S100A8/A9 (E) were quantified by ELISA. Representative data are shown. **(D, F)** Cells were stimulated with TNF (10 ng/mL) plus z-VAD-fmk (10 μM) in the presence or absence of Nec-1s (10 μM), GSK872 (10 μM), or NSA (10 μM) for 24 h. Extracellular HMGB1 (D) and S100A8/A9 (F) were quantified by ELISA. Representative data are shown. **(G)** Proposed model illustrating enhanced cell death and DAMP release in *UBA1*-mutated cells and their possible downstream inflammatory consequences.

Because TNF plus z-VAD-fmk induced prominent RIPK1–RIPK3–MLKL signaling and pathway-dependent membrane permeabilization (Figures 3A and 4B), we next examined whether this pathway also contributed to extracellular ATP accumulation. Pharmacological inhibition of RIPK1, RIPK3, or MLKL with Nec-1s, GSK872, or NSA, respectively, attenuated the increase in extracellular ATP under TNF plus z-VAD-fmk conditions (Figure 5B). These findings are consistent with a contribution of necroptotic signaling to ATP release under caspase-inhibited conditions.

We next examined whether this extracellular phenotype extended to other DAMPs. High mobility group box 1 (HMGB1) and S100A8/A9 levels were also higher in culture supernatants from *UBA1*-mutated cells than from WT cells, with the largest genotype-associated differences observed following TNF plus z-VAD-fmk treatment (Figure 5C, E). Under this condition, inhibition of RIPK1, RIPK3, or MLKL attenuated extracellular HMGB1 and S100A8/A9 levels (Figure 5D, F). Similar increases in HMGB1 and S100A8/A9 were observed in the bulk-derived series (Supplementary Figure S9B, C). Together, these findings indicate that enhanced cell death in *UBA1*-mutated cells is accompanied by increased extracellular accumulation of multiple DAMPs, particularly under conditions associated with necroptotic cell death.

We also examined whether the increased extracellular DAMP levels were accompanied by broadly increased inflammatory cytokine production. Differences in IL-1β, IL-6, and IL-8 concentrations between WT and *UBA1*-mutated cells were limited under the conditions examined. IL-1β and IL-6 remained low, whereas IL-8 levels were broadly comparable between genotypes (Supplementary Figures S9D-F and S10A-C). Thus, the extracellular phenotype observed in *UBA1*-mutated cells was more prominently characterized by increased DAMP levels than by increased production of the inflammatory cytokines examined.

As an exploratory analysis, we examined whether extracellular factors released from *UBA1*-mutated cells affected NF-κB signaling. Following neutralization of residual TNF with the TNF inhibitor etanercept, conditioned media from *UBA1*-mutated cells tended to induce higher NF-κB reporter activity than those from WT cells (Supplementary Figure S10D). Direct coculture of NF-κB reporter cells with either WT or *UBA1*-mutated U937 cells showed a similar tendency (Supplementary Figure S10E).

Overall, these findings show that enhanced cell death in *UBA1*-mutated cells is accompanied by increased extracellular release of ATP, HMGB1, and S100A8/A9, supporting DAMP release as an extracellular consequence of heightened cell-death susceptibility in this model.

## Discussion

In the present study, genome editing yielded maintainable *UBA1*-mutated U937 cells that recapitulated multiple molecular and cellular features associated with VEXAS syndrome. Using this system, we showed that UBA1 dysfunction was associated with a modest increase in basal cell death and heightened susceptibility to TNF-dependent cell death. The predominant mode of cell-death signaling varied according to the experimental context: apoptotic signaling was prominent following TNF stimulation alone or TNF plus LCL161, whereas RIPK1–RIPK3–MLKL-dependent necroptotic signaling became prominent when caspase activity was inhibited with z-VAD-fmk. We further showed that enhanced cell death was accompanied by increased extracellular accumulation of ATP, HMGB1, and S100A8/A9. Together, these findings define a cell-death-prone phenotype of *UBA1*-mutated myeloid cells and identify increased extracellular DAMP accumulation as a downstream consequence of this heightened cell-death susceptibility.

An unexpected but informative feature of model generation was that, despite extensive screening, we did not isolate a maintainable clone carrying the intended M41 mutation on both *UBA1* alleles without an accompanying on-target deletion. Although less common than small insertions or deletions, larger unintended deletions extending beyond the Cas9 cleavage site have been reported as on-target outcomes of CRISPR/Cas9 editing [19–21]. Thus, the deletion observed in our clones is consistent with a recognized, although relatively infrequent, outcome of genome editing. *UBA1* has been identified as an essential gene in multiple genome-wide loss-of-function screens [22–24], raising the possibility that strong UBA1 dysfunction imposed selective pressure during clonal expansion. The recovered clones instead carried the M41 mutation on one allele and a deletion on the other, suggesting that this allelic configuration may have permitted substantial UBA1 dysfunction while maintaining cellular viability. Transcripts from the deletion allele retained downstream exons, including exon 4 containing the alternative M67 translation initiation site, potentially allowing residual UBA1c expression. Although the functional significance of this transcript remains speculative, these findings suggest that UBA1 allelic configuration may influence the ability of cells with substantial UBA1 dysfunction to persist.

The resulting maintainable cell system has particular value when considered alongside existing experimental models of VEXAS syndrome. Genome-edited HSPC models, primary macrophage systems, and engineered myeloid cell lines have provided complementary insights into UBA1 dysfunction [10–14, 25]. Recent studies further indicate that the consequences of UBA1 dysfunction differ across hematopoietic compartments, with mutant HSPCs showing proteostatic stress and myeloid bias, whereas mutant macrophages and monocytes exhibit prominent abnormalities in inflammatory cell-death responses [10, 11]. Notably, a recent engineered THP-1 model used a Tet-On/Off system to maintain exogenous UBA1b expression during the establishment of UBA1-mutant clones; depletion of exogenous UBA1b under Tet-Off conditions was accompanied by reduced cell viability and proliferation [25]. This highlights a practical feature of our model: despite increased basal cell death, the *UBA1*-mutated U937 clones can be maintained under standard culture conditions without inducible UBA1b rescue. This maintainability permits repeated biochemical, real-time functional, and pharmacological analyses using genetically characterized clones. Moreover, the principal phenotypes observed in the single-clone-derived series were reproduced in the independently generated bulk-derived series, reducing the likelihood that the major findings were restricted to a particular clone. Thus, this system complements existing VEXAS models by enabling detailed analysis of the signaling and extracellular consequences of UBA1-associated cell-death susceptibility.

Our findings suggest that UBA1 dysfunction may place myeloid cells in a cell-death-prone state. Even in the absence of stimulation, *UBA1*-mutated cells showed increased basal abundance of RIPK1, RIPK3, and MLKL, accompanied by a modest increase in spontaneous cell death. Notably, the increased basal abundance of RIPK1, RIPK3, and MLKL proteins was not accompanied by consistent increases in their mRNA levels, suggesting that post-transcriptional regulation or altered protein turnover may contribute, although the underlying mechanism remains unknown. Previous studies have shown that the abundance of core necroptotic proteins, including RIPK1, RIPK3, and MLKL, can influence cellular competence for necroptosis [26–28]. The coordinated increase in these proteins may therefore increase cellular competence for necroptotic pathway engagement under permissive conditions. Importantly, elevated protein abundance does not itself indicate constitutive activation of necroptosis. More broadly, *UBA1*-mutated cells exhibited greater cell death across a range of TNF concentrations, including at low concentrations, suggesting a reduced threshold for TNF-induced cell death. Together, these findings support a model in which UBA1 dysfunction is associated with a death-prone myeloid state characterized by heightened sensitivity to TNF-dependent cell death rather than constitutive activation of a single death pathway.

Although *UBA1*-mutated cells were broadly sensitized to TNF-dependent cell death, the mode of cell-death execution depended on the signaling context. TNF stimulation alone or TNF plus LCL161 was associated predominantly with enhanced apoptotic signaling, as reflected by increased caspase activation and PARP1 cleavage. In contrast, when caspase activity was inhibited, RIPK1–RIPK3–MLKL signaling became prominent, and pharmacological inhibition of RIPK1, RIPK3, or MLKL markedly suppressed membrane permeabilization. These findings suggest that UBA1 dysfunction does not selectively prime a single death pathway but instead increases susceptibility to TNF-dependent cell death, with the downstream execution pathway determined by the signaling context.

Ubiquitin-dependent regulation of TNFR1 complex I provides a plausible mechanistic framework for this broader sensitization. Lys63- and Met1-linked ubiquitination of TNFR1 complex I contributes to stabilization of the complex and promotes pro-survival signaling, whereas disruption of these ubiquitin-dependent processes can favor formation of downstream death-inducing complexes [29–32]. Impaired Lys63-linked and linear polyubiquitination of RIPK1 has recently been demonstrated in *UBA1*-mutated macrophages [10]. Such impaired ubiquitin-dependent regulation could facilitate transition toward death-inducing signaling and may thereby enhance apoptotic signaling when caspase activity is intact and RIPK1–RIPK3–MLKL-dependent necroptosis when caspases are inhibited. Our observation that the basal abundance of several TNFR1-associated signaling proteins was altered is also consistent with broader remodeling of this signaling machinery, although the functional significance of these changes remains to be established. Notably, TNF-induced canonical NF-κB activation was broadly comparable between WT and *UBA1*-mutated cells, suggesting that UBA1 dysfunction does not uniformly enhance TNF signaling but may preferentially increase susceptibility of the cell-death branch.

Enhanced cell death in *UBA1*-mutated cells was accompanied by increased extracellular accumulation of ATP, HMGB1, and S100A8/A9. DAMPs released during regulated cell death can link cellular injury to inflammation [33, 34], and both HMGB1 and S100A8/A9 have established proinflammatory functions [35–37]. Notably, circulating DAMPs have also been reported to be elevated in patients with VEXAS syndrome [38], although their cellular sources and pathogenic significance remain unclear. In our model, the increase in extracellular DAMPs was more prominent than changes in the inflammatory cytokines examined: IL-1β and IL-6 remained low, whereas IL-8 levels were broadly comparable between WT and *UBA1*-mutated cells. This suggests that the extracellular phenotype associated with UBA1 dysfunction is not simply characterized by generalized cytokine production and raises the possibility that cell death-associated DAMP accumulation contributes to the inflammatory milieu in VEXAS syndrome. In exploratory conditioned-medium experiments, supernatants from *UBA1*-mutated cells showed a tendency to enhance NF-κB reporter activity after TNF neutralization. These observations raise the possibility that extracellular factors released in association with enhanced cell death may influence surrounding cells, although they do not establish a causal role for DAMPs in inflammatory activation.

Several limitations should be considered. First, U937 cells harbor two X chromosomes and therefore do not reproduce the typical hemizygous *UBA1*-mutant genotype of male VEXAS syndrome. Moreover, all long-term surviving clones carried the intended M41 mutation on one allele and an on-target deletion on the other; therefore, the observed phenotypes cannot be attributed solely to the M41 mutation. Second, the leukemic genetic and epigenetic background of U937 cells may modify the cellular consequences of UBA1 dysfunction. Third, the limited number of independently derived clones precluded formal genotype-level statistical comparisons. Finally, validation in primary patient-derived cells will be required to establish the physiological relevance and generalizability of these findings.

In conclusion, we characterized a maintainable *UBA1*-mutated human myeloid model that recapitulates multiple molecular and cellular features associated with VEXAS syndrome. Using this model, we showed that UBA1 dysfunction is associated with a cell-death-prone state with heightened susceptibility to TNF-dependent apoptotic and necroptotic cell death, accompanied by increased extracellular DAMP accumulation. These findings provide insight into how UBA1 dysfunction alters myeloid cell-death responses and their extracellular consequences and establish a tractable platform for further mechanistic studies of VEXAS syndrome.

## List of abbreviations

ATP: adenosine triphosphate
cIAP: cellular inhibitor of apoptosis protein
DAMP: damage-associated molecular pattern
ELISA: enzyme-linked immunosorbent assay
FADD: Fas-associated death domain protein
GFP: green fluorescent protein
HMGB1: high mobility group box 1
HSPC: hematopoietic stem/progenitor cell
LDH: lactate dehydrogenase
MLKL: mixed lineage kinase domain-like protein
NF-κB: nuclear factor kappa B
PARP1: poly(ADP-ribose) polymerase 1
PCR: polymerase chain reaction
RIPK1: receptor-interacting protein kinase 1
RIPK3: receptor-interacting protein kinase 3
RT-PCR: reverse-transcription polymerase chain reaction
TNF: tumor necrosis factor
TNFR1: tumor necrosis factor receptor 1
TRADD: TNFR1-associated death domain protein
VEXAS: vacuoles, E1 enzyme, X-linked, autoinflammatory, somatic
WT: wild-type

## Declarations

### Ethics approval and consent to participate

All animal experiments were approved by the Institutional Animal Care and Use Committee of Shigei Medical Research Institute (approval number: 24002). All experimental procedures were conducted according to institutional and NIH guidelines for the humane use of animals.

### Consent for publication

Not applicable.

### Availability of data and materials

The datasets generated and/or analyzed during the current study are available from the corresponding author on reasonable request.

### Competing interests

YSa received a research grant from Nippon Shinyaku Co., Ltd. The remaining authors declare that they have no competing interests.

### Funding

This work was supported by JSPS KAKENHI [24K11570 to YSa; 26K11352 to NB; 24K02491 to TM], a Health Labor Sciences Research Grant [23FC1016 and 26FC1017 to TM], Drug Discovery Booster Program of the Japan Agency for Medical Research and Development (AMED) (DNW-26013 to TM), Promotion and Mutual Aid Corporation for Private Schools of Japan (TM), Okayama Prefecture Special Power-Source Prefecture Science and Technology Promotion Grant (TM), The Japan College of Rheumatology (JCR) Grant for Promoting Research for D2T RA (TM), The JCR Research Grant for Optimizing Medical Care for Late-Onset Rheumatoid Arthritis (TM), Kawasaki Medical School Project Grants (R07B-066 to YSa, R07B-067 to NB, R07B-064 to TM), the Ryobi Teien Memorial Foundation (to YSa), the Teraoka Memorial Scholarship Foundation (to YSa), the Kanahara Ichiro Memorial Foundation for Medicine and Medical Care (to YSa), the Okayama Medical Foundation (to YSa), the Japan Rheumatism Foundation (to YSa), Nippon Shinyaku Co., Ltd. Research Grant (to YSa), the Kurozumi Medical Foundation (to YSa), the Yokoyama Foundation for Clinical Pharmacology (to YSa), the Wesco Scientific Promotion Foundation (to YSa), and the Foundation for Advanced Research in Medical Science (to YSa).

### Authors’ contributions

YSa and TM conceived and designed the study. YSa performed the majority of the experiments and drafted the manuscript. NB and MI performed experiments. EK performed genetic analyses. MM and TK generated monoclonal antibodies. YSa, NB, MI, EK, MM, TK, YSh, SM, TI, and TM analyzed and interpreted the data. TM supervised the study and acquired funding. All authors critically reviewed the manuscript and approved the final version.

## Acknowledgments

We thank Keiko Watanabe, Noriko Miyake, Hiroki Usui, Haruko Ohta, Minami Shimizu (Department of Immunology and Molecular Genetics, Kawasaki Medical School); Yoshinori Matsumoto, Takayuki Katsuyama, Haruki Watanabe (Department of Nephrology, Rheumatology, Endocrinology and Metabolism, Okayama University Faculty of Medicine, Dentistry and Pharmaceutical Sciences); Shin Morizane, Yoshihiro Matsuda, Daiki Takezaki (Department of Dermatology, Okayama University Faculty of Medicine, Dentistry and Pharmaceutical Sciences); Shuichiro Okamoto (Department of Biochemistry, Kawasaki Medical School); Natsumi Okazaki, Sayaka Uji (Kawasaki University of Medical Welfare) for their technical assistance, and Yasuhiko Hayakawa (Nepa Gene Co., Ltd., Ichikawa, Chiba, Japan) for his support in optimizing electroporation. We are also grateful to the staff of the Central Research Institute of Kawasaki Medical School.

The authors used AI-based language tools, ChatGPT (OpenAI) and Gemini (Google), for grammar checking and improving the readability of the manuscript. The authors are solely responsible for the scientific content.

## Supplementary Figure legends

**Supplementary Figure S1. Molecular characterization of single-clone-derived WT and *UBA1*-mutated U937 cells. (A)** Schematic representation of primer locations used for genomic DNA analysis (upper) and transcript structural analysis (lower) of the *UBA1* gene. **(B)** Representative agarose gel images showing PCR products generated using the primer sets illustrated in (A). **(C)** Long-read sequencing read coverage across the *UBA1* locus obtained using the PromethION platform. **(D)** Relative *UBA1* transcript expression measured by quantitative PCR using three primer pairs targeting the exon 1–2 junction/exon 2, exons 5–6, and the exon 15–16 junction/exon 16. Data are presented as mean ± SD of three independent experiments.

**Supplementary Figure S2. Characterization of the bulk-derived *UBA1*-mutated U937 cell series.** Bulk-derived U937 cells carrying *UBA1* M41 mutations were established independently. An independently established WT clone (WT #10) was included as an additional WT control throughout the analyses. **(A)** Genomic PCR and RT-PCR analyses of the bulk-derived WT and *UBA1*-mutated cells using the primer sets shown in Supplementary Figure S1. Representative agarose gel images are shown. **(B)** Schematic summary of the genomic and transcript structures of *UBA1* in the bulk-derived WT and *UBA1*-mutated cells. Genomic structures of the two *UBA1* alleles are shown on the left, and the corresponding transcript structures are shown on the right. Reduced expression of transcripts containing exons 2 and 3 in the *UBA1*-mutated cells is indicated. **(C)** IGV visualization of long-read sequencing read coverage across the *UBA1* locus corresponding to the genomic alterations shown in (B). **(D)** Immunoblot analysis of UBA1 isoform expression in bulk-derived WT and *UBA1*-mutant clones. The upper antibody recognizes UBA1a, UBA1b, and UBA1c, whereas the lower antibody recognizes UBA1a and UBA1b. Representative immunoblots are shown. **(E)** Cell proliferation was assessed using the CCK-8 assay. Representative data from triplicate wells are shown. **(F)** Cell viability (blue) and membrane permeabilization (red) were monitored under basal culture conditions using the RealTime-Glo™ MT Cell Viability Assay and CellTox™ Green Cytotoxicity Assay. Representative data from duplicate wells are shown. WT represents the combined results of WT #7 and WT #10. Data from the M41V #243 cells are not shown because the corresponding experiment was not performed. **(G)** Cell-cycle distribution determined by flow cytometry.

**Supplementary Figure S3. Long-read sequencing analysis of the single-clone-derived WT and *UBA1*-mutated U937 cells.** Long-range PCR products generated using four primer pairs spanning approximately 20 kb, 5 kb, 2 kb, and 600 bp around the *UBA1* M41 region were subjected to PromethION sequencing. Integrative Genomics Viewer (IGV) views of the aligned sequencing reads are shown for WT #7 **(A)**, M41V #52 **(B)**, and M41V #164 **(C)**. For each clone, the upper panel shows an overview of the *UBA1* locus, and the lower panel shows the M41 region at higher magnification.

**Supplementary Figure S4. Long-read sequencing analysis of the bulk-derived WT #10 and *UBA1*-mutated U937 cells.** Long-range PCR products generated using four primer pairs spanning approximately 20 kb, 5 kb, 2 kb, and 600 bp around the *UBA1* M41 region were subjected to PromethION sequencing. IGV views of the aligned sequencing reads are shown for WT #10 **(A)**, M41V #243 **(B)**, M41T #15 **(C)**, and M41L #132 **(D)**. For each clone, the upper panel shows an overview of the *UBA1* locus, and the lower panel shows the M41 region at higher magnification.

**Supplementary Figure S5. Cell death phenotypes of the bulk-derived *UBA1*-mutated U937 cell series.** Bulk-derived WT and *UBA1*-mutated U937 cells were analyzed. **(A)** Spontaneous LDH release under basal culture conditions was measured for the indicated times. Representative data from triplicate wells are shown. **(B)** Cell death under basal culture conditions was analyzed by Annexin V/PI staining after 24 h. Representative flow cytometry plots are shown below. Data are presented as mean ± SD of two independent experiments. **(C)** Cells were stimulated with the indicated concentrations of TNF for 24 h, and LDH release was measured. The upper panel shows LDH release expressed as a percentage of maximum LDH release, and the lower panel shows ΔLDH release relative to untreated cells (TNF 0 ng/mL). Representative data from triplicate wells are shown. WT represents the combined results of WT #7 and WT #10.

**Supplementary Figure S6. Additional analysis of TNF signaling in single-clone-derived *UBA1*-mutated cells. (A)** WT and M41V U937 cells were stimulated with TNF (10 ng/mL) in the presence of LCL161 (250 nM) for the indicated times. Cell lysates were analyzed by immunoblotting for caspase-3, cleaved caspase-3, and PARP1. β-actin served as a loading control. Full-length and cleaved caspase-3 detected with the same antibody are shown separately because different exposure times were used to visualize the cleaved form. **(B)** WT and M41V U937 cells were stimulated with TNF (10 ng/mL) for the indicated times. Cell lysates were analyzed by immunoblotting for phosphorylated and total IκB, phosphorylated and total NF-κB p65, and NF-κB p105/p50. β-actin served as a loading control. **(C)** Immunoblot analysis of TNFR1 signaling components and related regulatory proteins under basal culture conditions. A20, cIAP1, HOIP, SHARPIN, OTULIN, TRADD, and FADD were analyzed as proteins involved in TNFR1 signaling. Representative immunoblots are shown in (A–C).

**Supplementary Figure S7. Cell death kinetics of WT and *UBA1*-mutated cells under different stimulation conditions.** Annexin V luminescence (blue) and membrane permeabilization-associated fluorescence (red), reflecting phosphatidylserine exposure and membrane permeabilization, respectively, were monitored using the RealTime-Glo™ Annexin V Apoptosis and Necrosis Assay. WT and M41V cells were left untreated or stimulated with TNF (10 ng/mL), TNF plus z-VAD-fmk (10 μM), TNF plus LCL161 (250 nM), or TNF plus LCL161 (250 nM) and z-VAD-fmk (10 μM) for 12 h.

**Supplementary Figure S8. Cell death kinetics of bulk-derived *UBA1*-mutated U937 cells.** Annexin V luminescence (blue) and membrane permeabilization-associated fluorescence (red), reflecting phosphatidylserine exposure and membrane permeabilization, respectively, were monitored using the RealTime-Glo™ Annexin V Apoptosis and Necrosis Assay. WT, M41T #15, and M41L #132 cells were left untreated or stimulated with z-VAD-fmk (10 μM), TNF (10 ng/mL), or TNF plus z-VAD-fmk (10 μM) for 24 h. WT represents the combined results of WT #7 and WT #10. Representative data are shown. Data from the M41V #243 cells are not shown because the corresponding experiment was not performed.

**Supplementary Figure S9. DAMP release and inflammatory cytokine levels in bulk-derived *UBA1*-mutated U937 cells. (A)** Extracellular ATP levels were measured at the indicated time points over 12 h under non-treatment, z-VAD-fmk (10 μM), TNF (10 ng/mL), or TNF (10 ng/mL) plus z-VAD-fmk (10 μM) conditions. Representative data from duplicate wells are shown. **(B, C)** Cells were stimulated with z-VAD-fmk (10 μM), TNF (10 ng/mL), or TNF (10 ng/mL) plus z-VAD-fmk (10 μM) for 24 h. Extracellular HMGB1 (B) and S100A8/A9 (C) were quantified by ELISA. Representative data are shown. **(D-F)** Cells were stimulated with z-VAD-fmk (10 μM), TNF (10 ng/mL), or TNF (10 ng/mL) plus z-VAD-fmk (10 μM) for 24 h. IL-1β (D), IL-6 (E), and IL-8 (F) concentrations in the culture supernatants were measured by ELISA. Representative data are shown. WT represents the combined results of WT #7 and WT #10. Data from the M41V #243 cells are not shown because the corresponding experiment was not performed.

**Supplementary Figure S10. Inflammatory cytokine levels and NF-**κ**B reporter activity in WT and *UBA1*-mutated cells. (A–C)** Cells were left untreated or stimulated with z-VAD-fmk (10 μM), TNF (10 ng/mL), or TNF (10 ng/mL) plus z-VAD-fmk (10 μM) for 24 h. IL-1β **(A)**, IL-6 **(B)**, and IL-8 **(C)** concentrations in the culture supernatants were measured by ELISA. Representative data are shown. **(D)** WT and *UBA1*-mutated U937 cells were stimulated with TNF (10 ng/mL) or TNF (10 ng/mL) plus z-VAD-fmk (10 μM), and culture supernatants were collected as conditioned media, as illustrated in the left panel. Residual TNF activity in the conditioned media was neutralized with etanercept before application to U937 NF-κB luciferase reporter cells. The middle and right panels show NF-κB reporter activity obtained using the single-clone-derived and independently established bulk-derived series, respectively. NF-κB reporter activity was measured as relative light units. Representative data from duplicate wells are shown. **(E)** NF-κB luciferase reporter cells were co-cultured in direct contact with WT or M41V #52 U937 cells for 48 or 96 h, and NF-κB reporter activity was measured as relative light units. Representative data from triplicate wells are shown.

## Notes

### Competing Interest Statement

YSa received a research grant from Nippon Shinyaku Co. Ltd. The remaining authors declare that the research was conducted in the absence of commercial or financial relationships that could be construed as potential conflicts of interest.

