## Supplementary Figures for "Characterization of a maintainable human myeloid model of VEXAS syndrome with enhanced TNF-induced cell death and DAMP release": Supplementary Figure 20260907_1900 Letter.pptx

### Slide 1
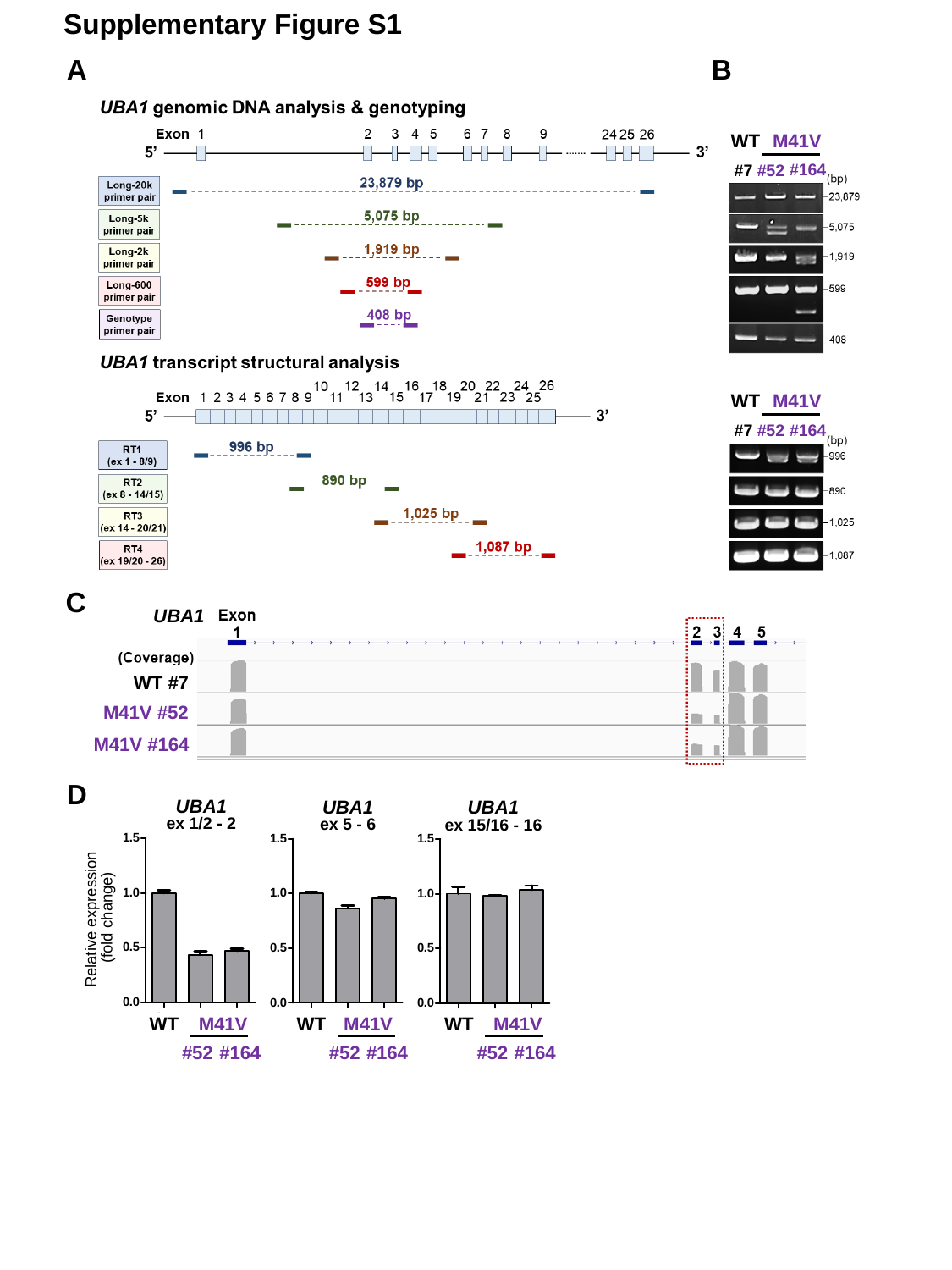

Supplementary Figure S1
A
B
WT
M41V
#164
#7
#52
WT
M41V
#164
#7
#52
C
UBA1
WT #7
M41V #52
M41V #164
D
UBA1
ex 1/2 - 2
UBA1
ex 5 - 6
UBA1
ex 15/16 - 16
Relative expression
 (fold change)
WT
M41V
WT
M41V
WT
M41V
#52
#52
#52
#164
#164
#164

### Slide 2
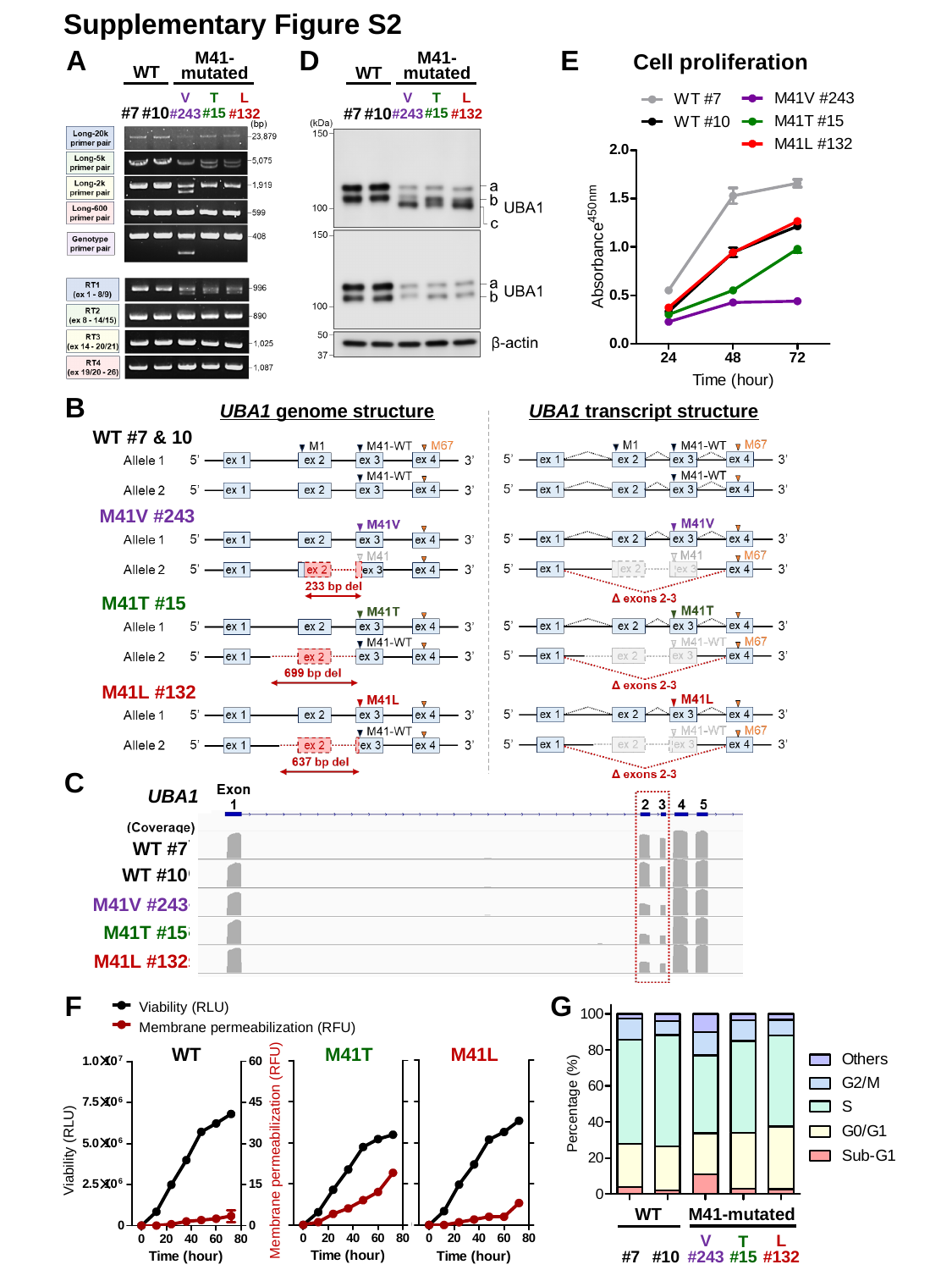

Supplementary Figure S2
D
A
E
Cell proliferation
M41-mutated
M41-mutated
WT
WT
T
#15
T
#15
V
#243
L
#132
V
#243
L
#132
#7
#10
#7
#10
B
UBA1 genome structure
UBA1 transcript structure
WT #7 & 10
M41V #243
M41T #15
M41L #132
C
UBA1
WT #7
WT #10
M41V #243
M41T #15
M41L #132
F
G
Viability (RLU)
Membrane permeabilization (RFU)
M41L
WT
M41T
Percentage (%)
Viability (RLU)
Membrane permeabilization (RFU)
M41-mutated
WT
L
#132
V
#243
T
#15
#7
#10

### Slide 3
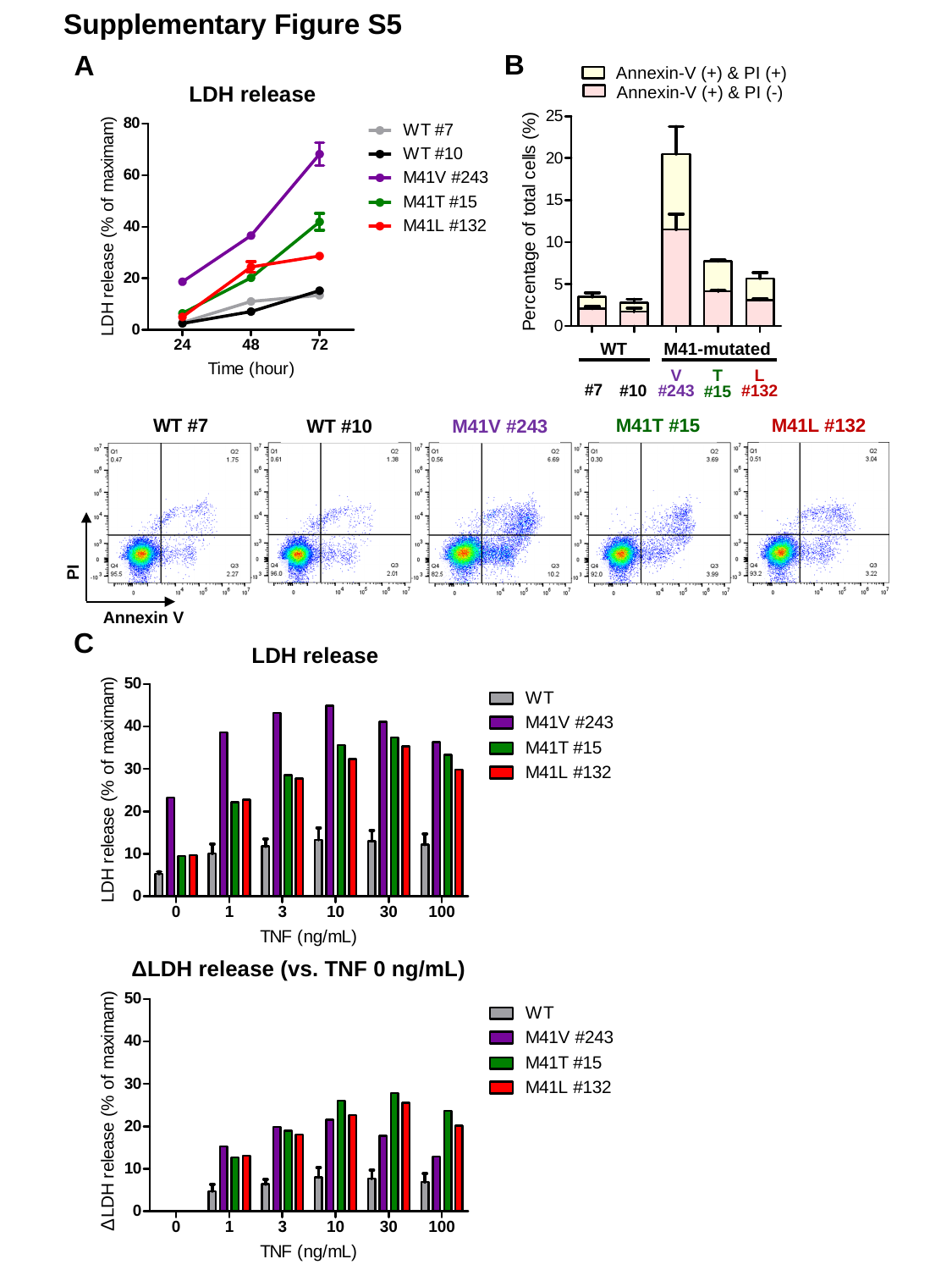

Supplementary Figure S5
B
A
Annexin-V (+) & PI (+)
LDH release
Annexin-V (+) & PI (-)
WT
M41-mutated
V
#243
L
#132
T
#15
#7
#10
M41T #15
M41L #132
WT #7
WT #10
M41V #243
PI
Annexin V
C
LDH release
Δ
ΔLDH release (vs. TNF 0 ng/mL)
Δ

### Slide 4
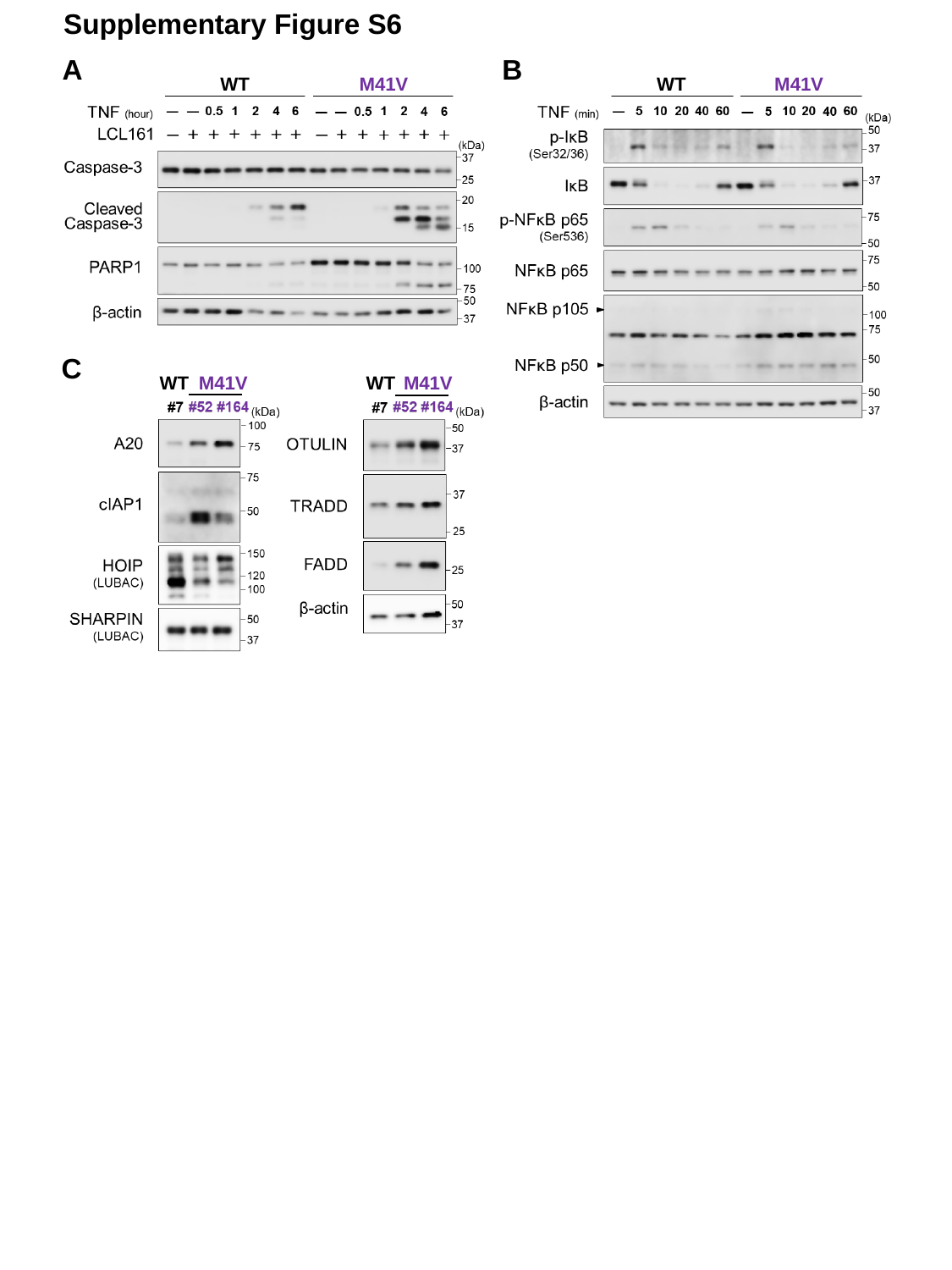

Supplementary Figure S6
A
B
M41V
M41V
WT
WT
C
M41V
WT
M41V
WT

### Slide 5
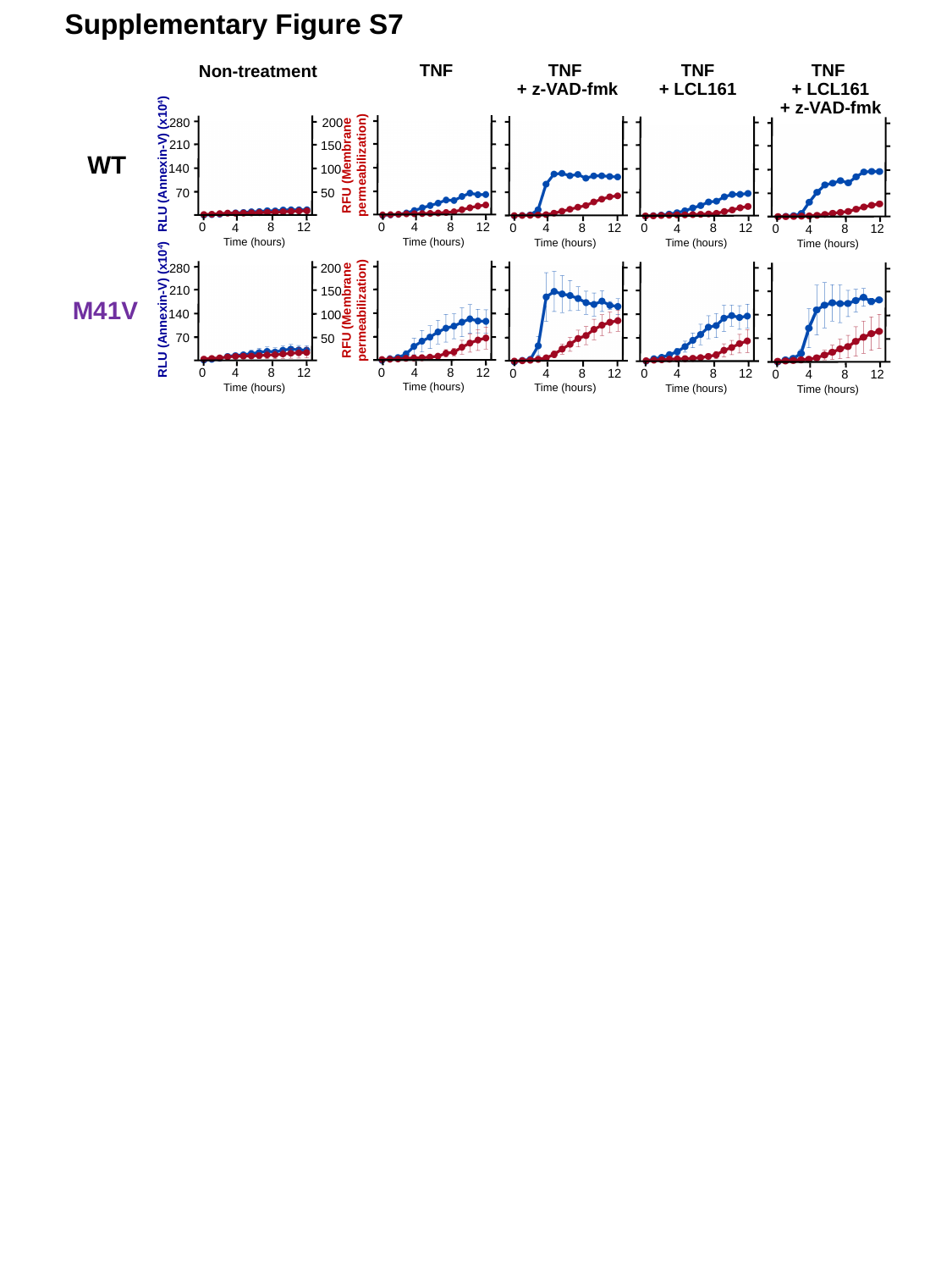

Supplementary Figure S7
Non-treatment
TNF
+ z-VAD-fmk
TNF
+ LCL161
TNF
TNF
+ LCL161
+ z-VAD-fmk
280
200
210
150
WT
RFU (Membrane
permeabilization)
RLU (Annexin-V) (x10^4^)
140
100
70
50
8
12
0
4
8
12
0
4
8
12
0
4
8
12
0
4
8
12
0
4
Time (hours)
Time (hours)
Time (hours)
Time (hours)
Time (hours)
280
200
210
150
M41V
RFU (Membrane
permeabilization)
RLU (Annexin-V) (x10^4^)
140
100
70
50
8
12
0
4
8
12
0
4
8
12
0
4
8
12
0
4
8
12
0
4
Time (hours)
Time (hours)
Time (hours)
Time (hours)
Time (hours)

### Slide 6
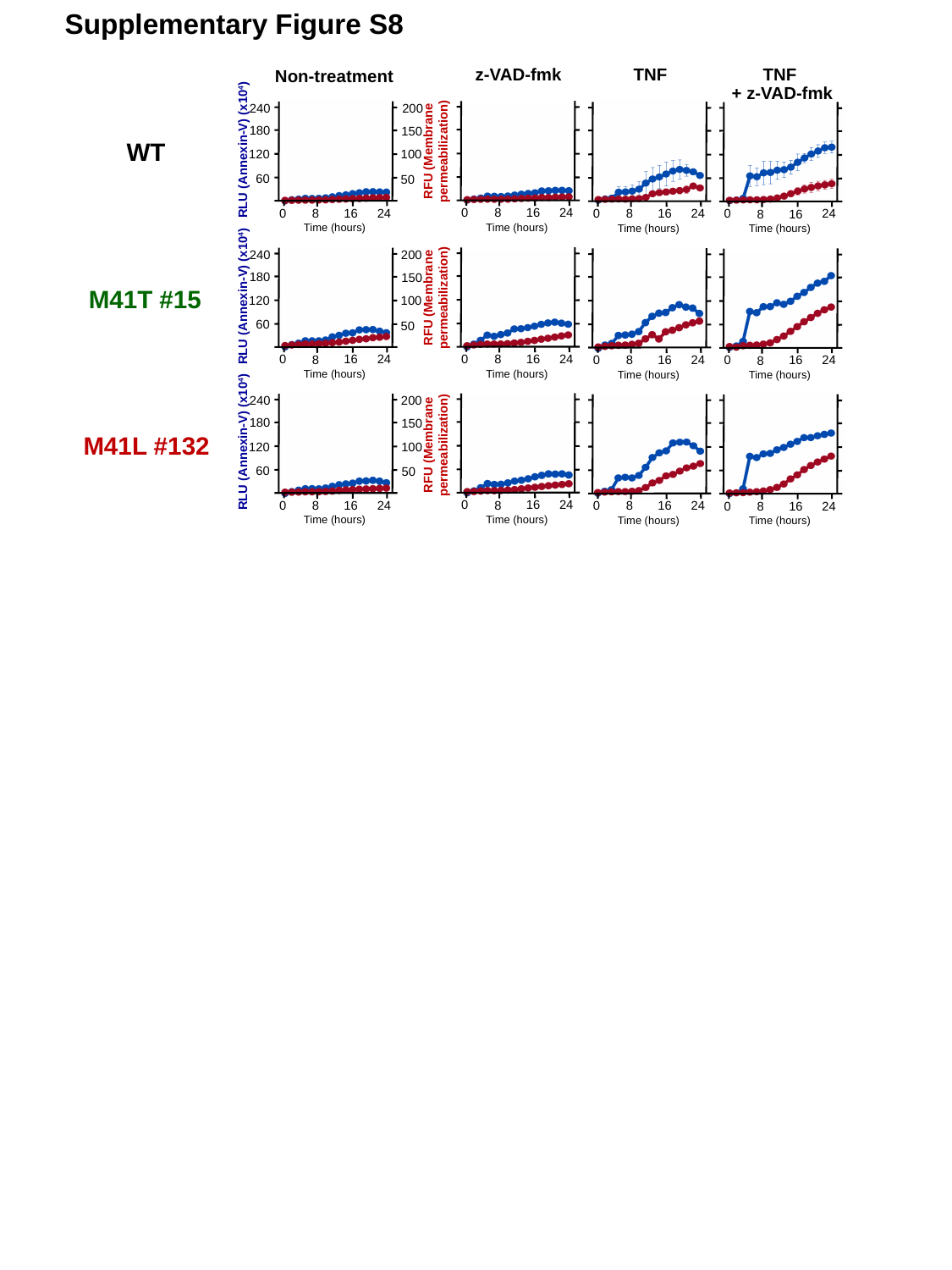

Supplementary Figure S8
Non-treatment
TNF
+ z-VAD-fmk
z-VAD-fmk
TNF
240
200
180
150
RFU (Membrane
permeabilization)
WT
RLU (Annexin-V) (x10^4^)
120
100
60
50
24
0
16
8
24
0
16
8
24
0
16
8
24
0
16
8
Time (hours)
Time (hours)
Time (hours)
Time (hours)
240
200
180
150
RFU (Membrane
permeabilization)
M41T #15
RLU (Annexin-V) (x10^4^)
120
100
60
50
24
0
16
8
24
0
16
8
24
0
16
8
24
0
16
8
Time (hours)
Time (hours)
Time (hours)
Time (hours)
240
200
180
150
RFU (Membrane
permeabilization)
M41L #132
RLU (Annexin-V) (x10^4^)
120
100
60
50
24
0
16
8
24
0
16
8
24
0
16
8
24
0
16
8
Time (hours)
Time (hours)
Time (hours)
Time (hours)

### Slide 7
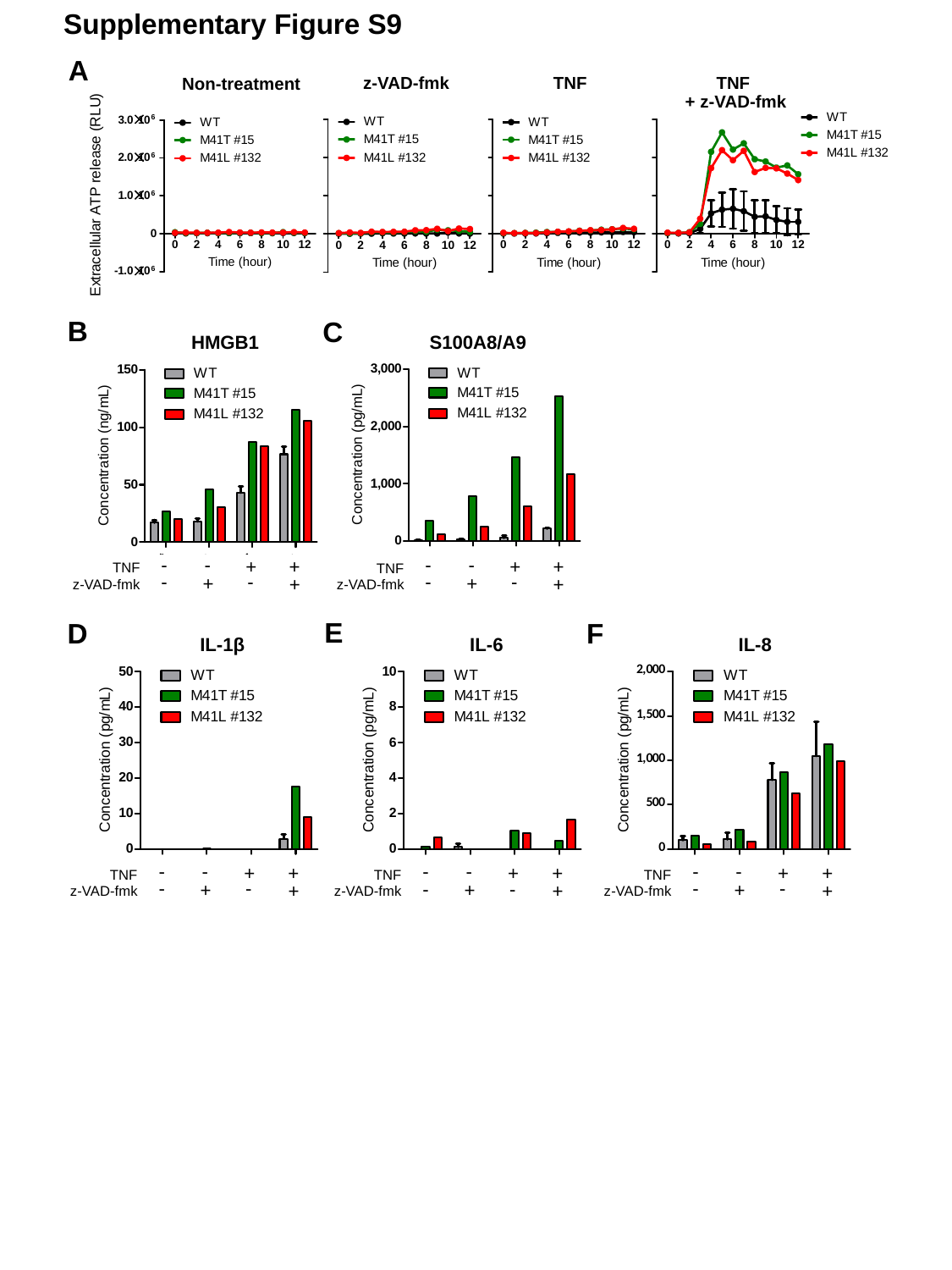

Supplementary Figure S9
A
Non-treatment
TNF
z-VAD-fmk
TNF
+ z-VAD-fmk
B
C
S100A8/A9
HMGB1
-
-
-
-
+
+
+
+
TNF
TNF
-
-
-
-
+
+
+
+
z-VAD-fmk
z-VAD-fmk
E
D
F
IL-1β
IL-6
IL-8
-
-
-
-
-
-
+
+
+
+
+
+
TNF
TNF
TNF
-
-
-
-
-
-
+
+
+
+
+
+
z-VAD-fmk
z-VAD-fmk
z-VAD-fmk

### Slide 8
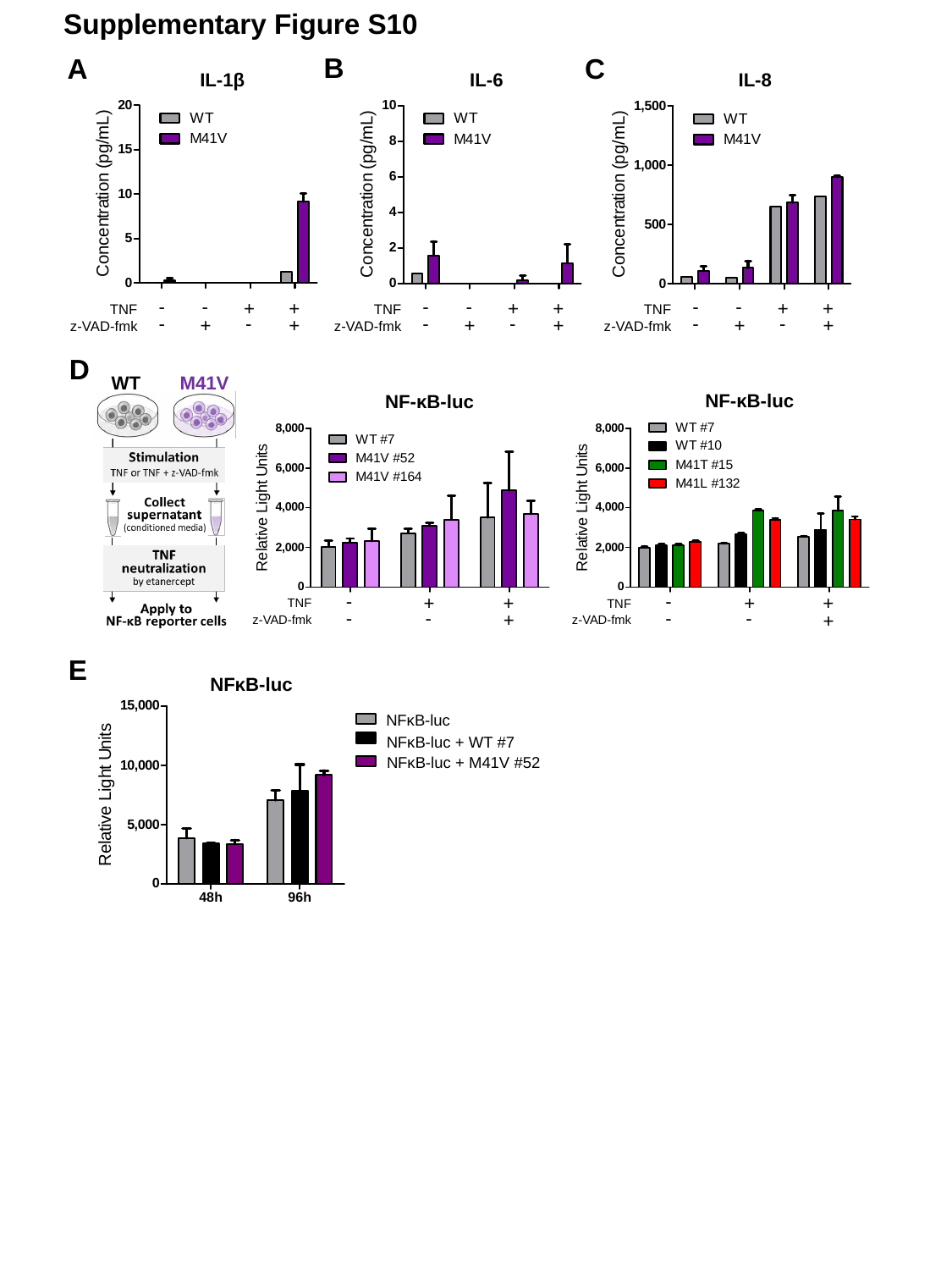

Supplementary Figure S10
B
A
C
IL-1β
IL-6
IL-8
-
-
-
-
-
-
+
+
+
+
+
+
TNF
TNF
TNF
-
-
-
-
-
-
+
+
+
+
+
+
z-VAD-fmk
z-VAD-fmk
z-VAD-fmk
D
WT
M41V
NF-κB-luc
NF-κB-luc
-
-
+
+
+
+
TNF
TNF
-
-
-
-
+
+
z-VAD-fmk
z-VAD-fmk
E
NFκB-luc
NFκB-luc
NFκB-luc + WT #7
NFκB-luc + M41V #52
