## Supplementary Figures for "Characterization of a maintainable human myeloid model of VEXAS syndrome with enhanced TNF-induced cell death and DAMP release": Supplementary Figure S3,S4 20260829 Letter.pptx

### Slide 1
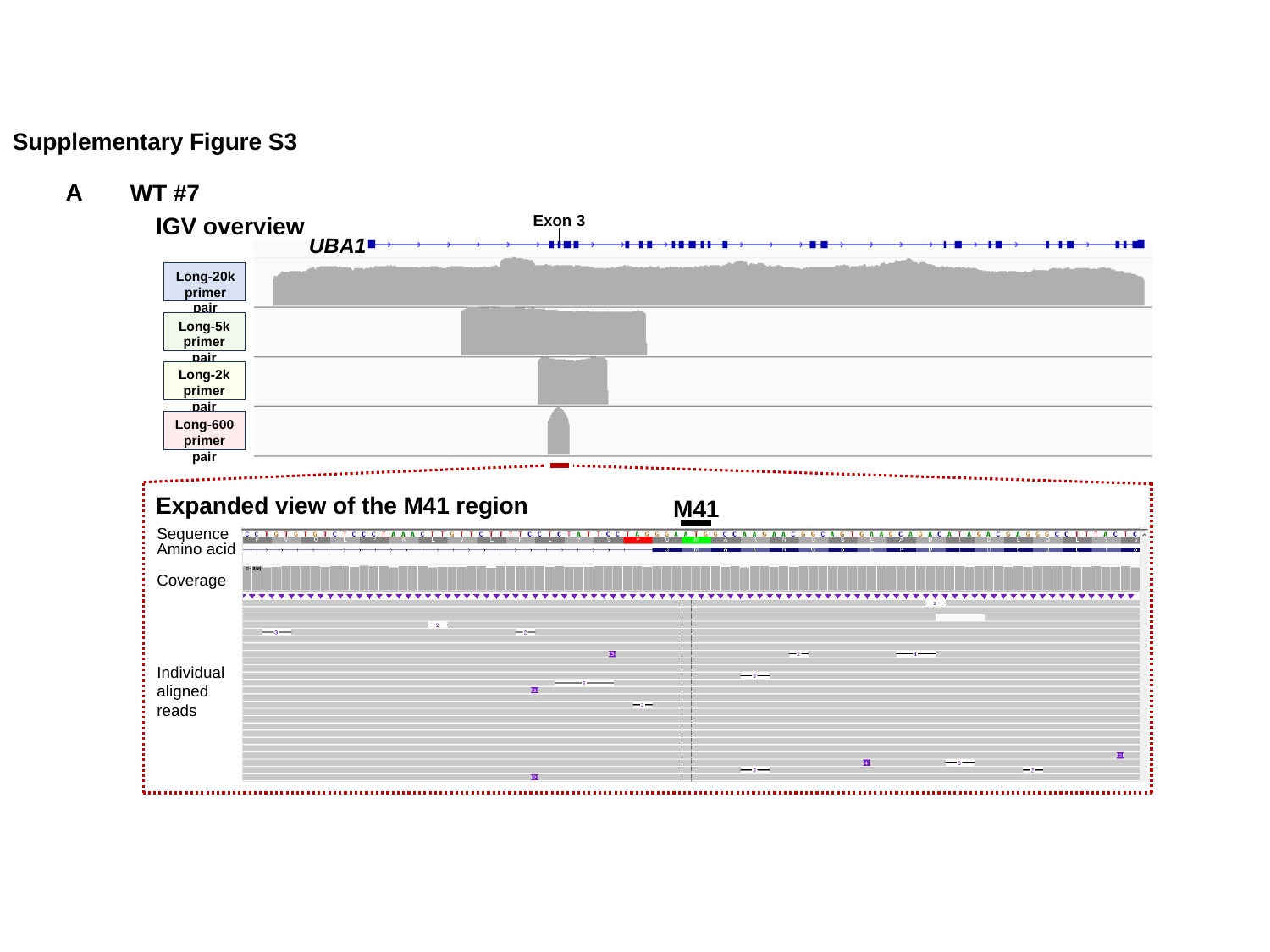

Supplementary Figure S3
A
WT #7
Exon 3
IGV overview
UBA1
Long-20k
primer pair
Long-5k
primer pair
Long-2k
primer pair
Long-600
primer pair
Expanded view of the M41 region
M41
Sequence
Amino acid
Coverage
Individual
aligned reads

### Slide 2
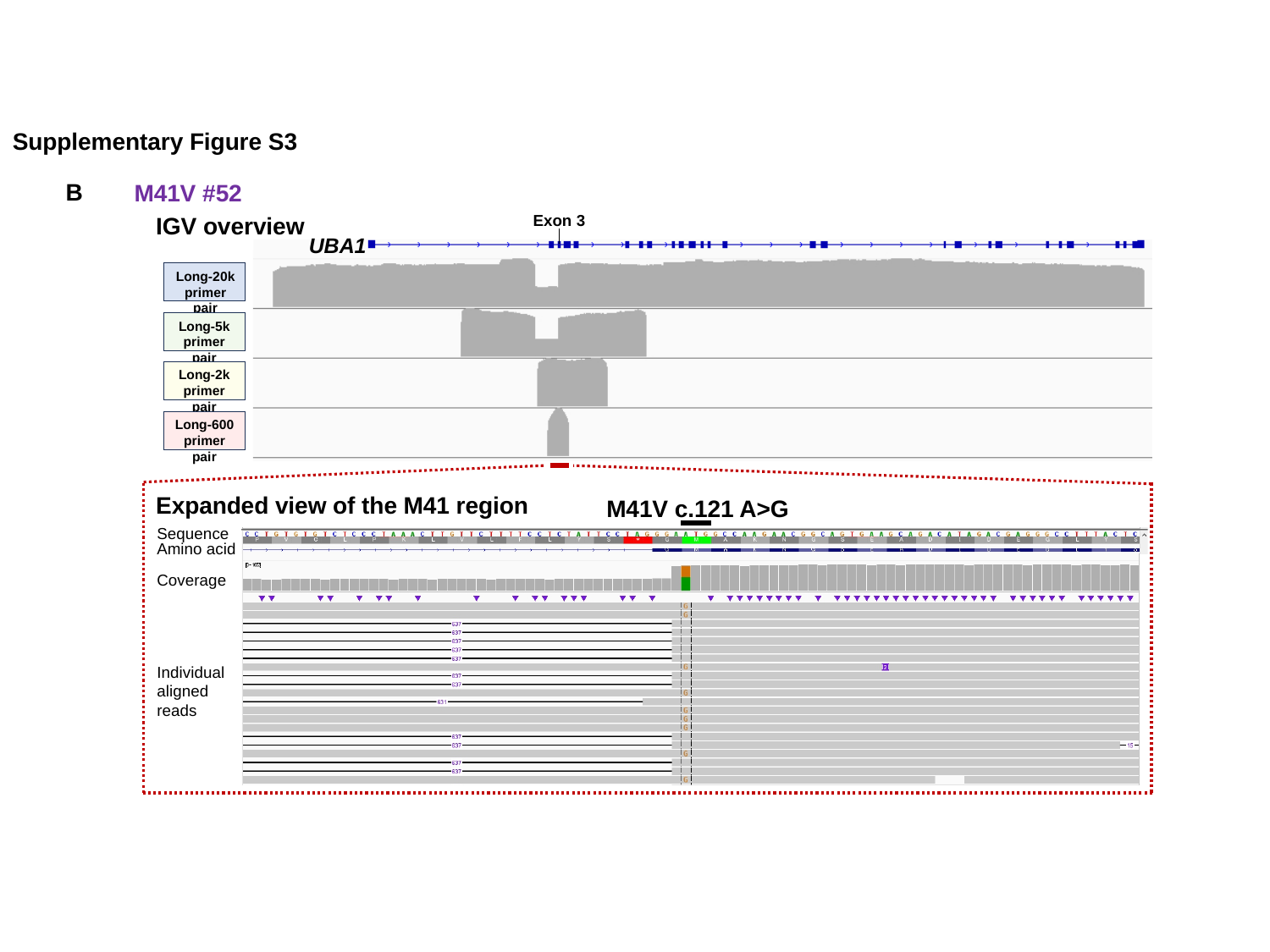

Supplementary Figure S3
B
M41V #52
Exon 3
IGV overview
UBA1
Long-20k
primer pair
Long-5k
primer pair
Long-2k
primer pair
Long-600
primer pair
Expanded view of the M41 region
M41V c.121 A>G
Sequence
Amino acid
Coverage
Individual
aligned reads

### Slide 3
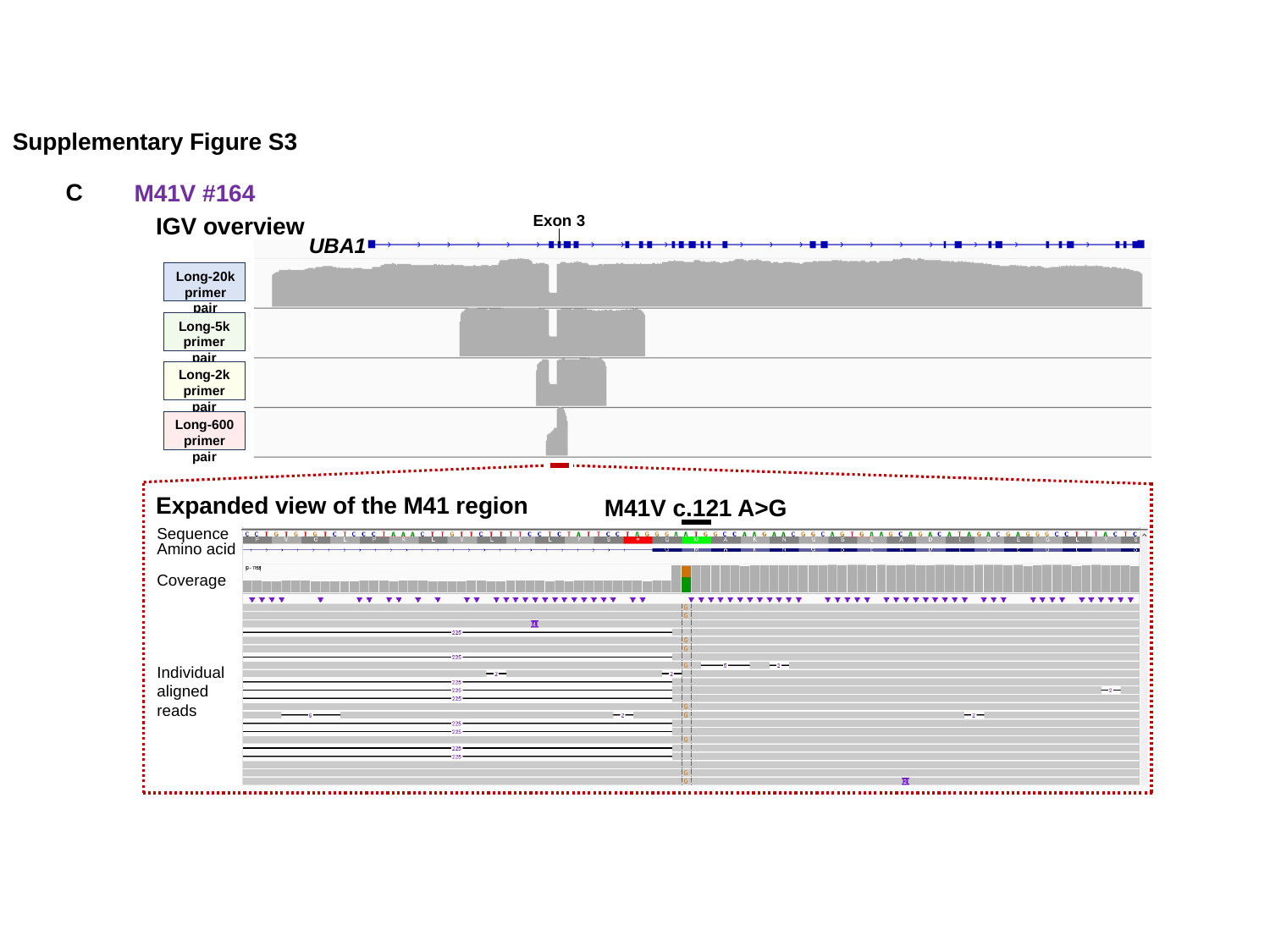

Supplementary Figure S3
C
M41V #164
Exon 3
IGV overview
UBA1
Long-20k
primer pair
Long-5k
primer pair
Long-2k
primer pair
Long-600
primer pair
Expanded view of the M41 region
M41V c.121 A>G
Sequence
Amino acid
Coverage
Individual
aligned reads

### Slide 4
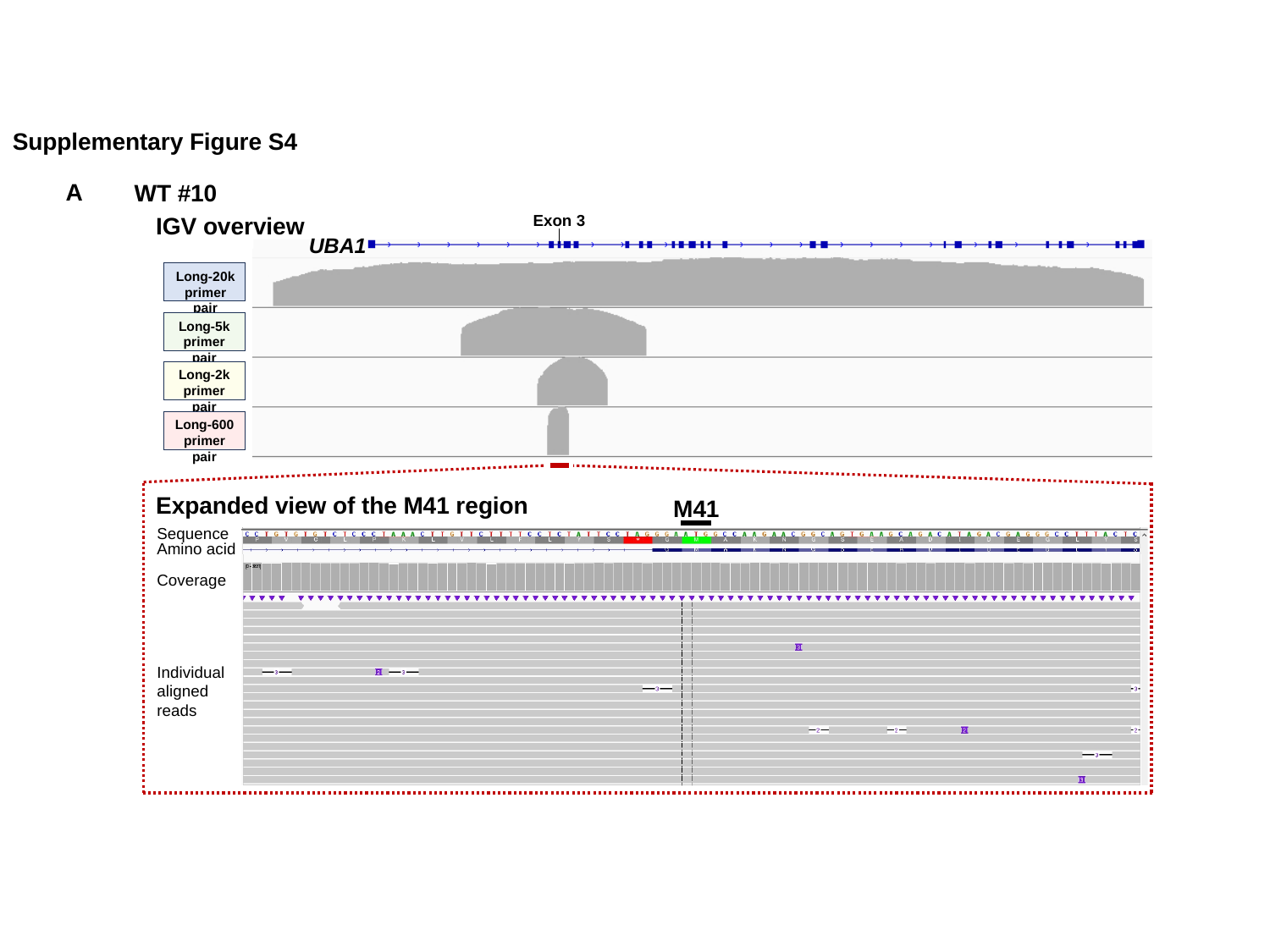

Supplementary Figure S4
A
WT #10
Exon 3
IGV overview
UBA1
Long-20k
primer pair
Long-5k
primer pair
Long-2k
primer pair
Long-600
primer pair
Expanded view of the M41 region
M41
Sequence
Amino acid
Coverage
Individual
aligned reads

### Slide 5
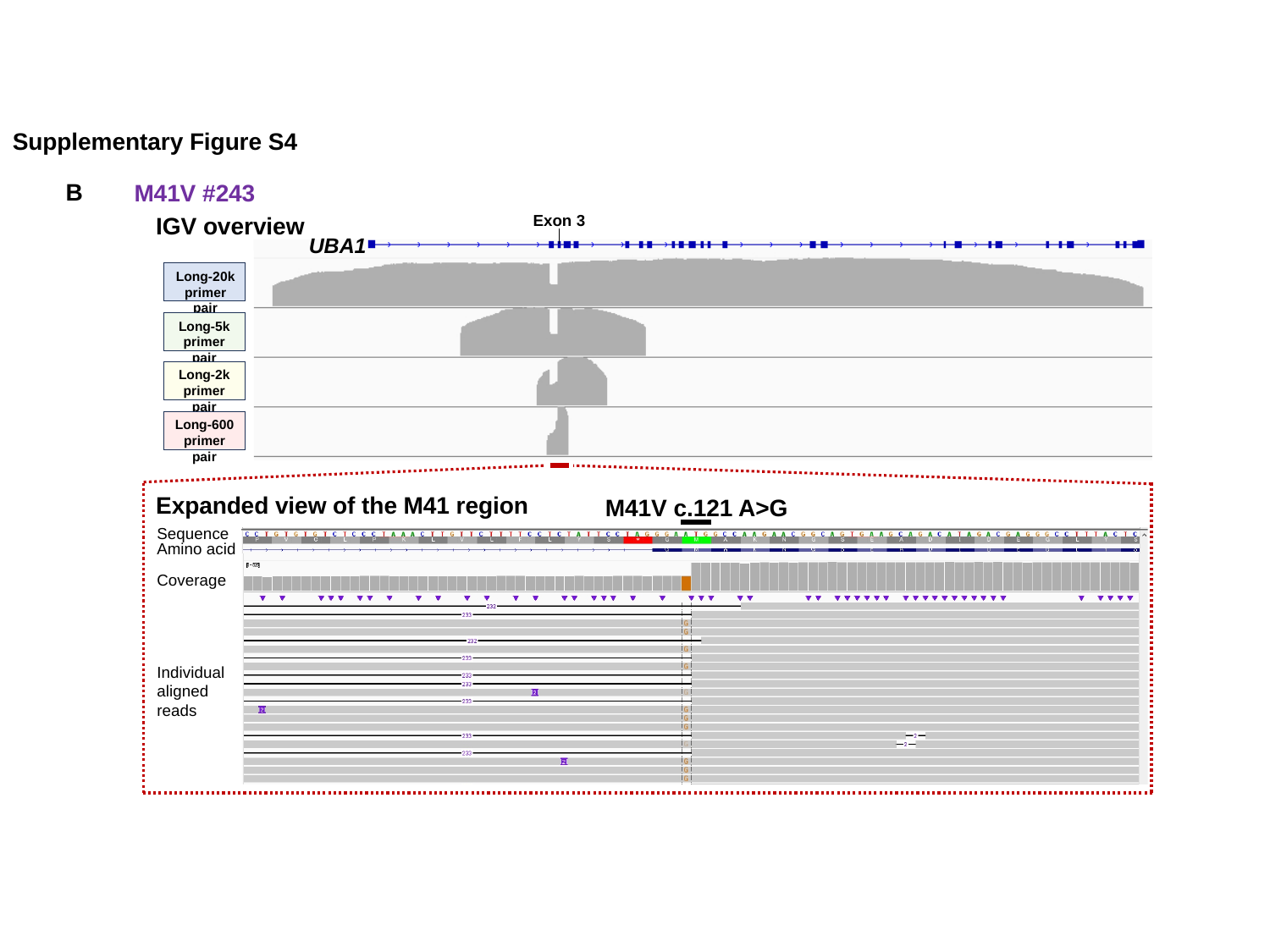

Supplementary Figure S4
B
M41V #243
Exon 3
IGV overview
UBA1
Long-20k
primer pair
Long-5k
primer pair
Long-2k
primer pair
Long-600
primer pair
Expanded view of the M41 region
M41V c.121 A>G
Sequence
Amino acid
Coverage
Individual
aligned reads

### Slide 6
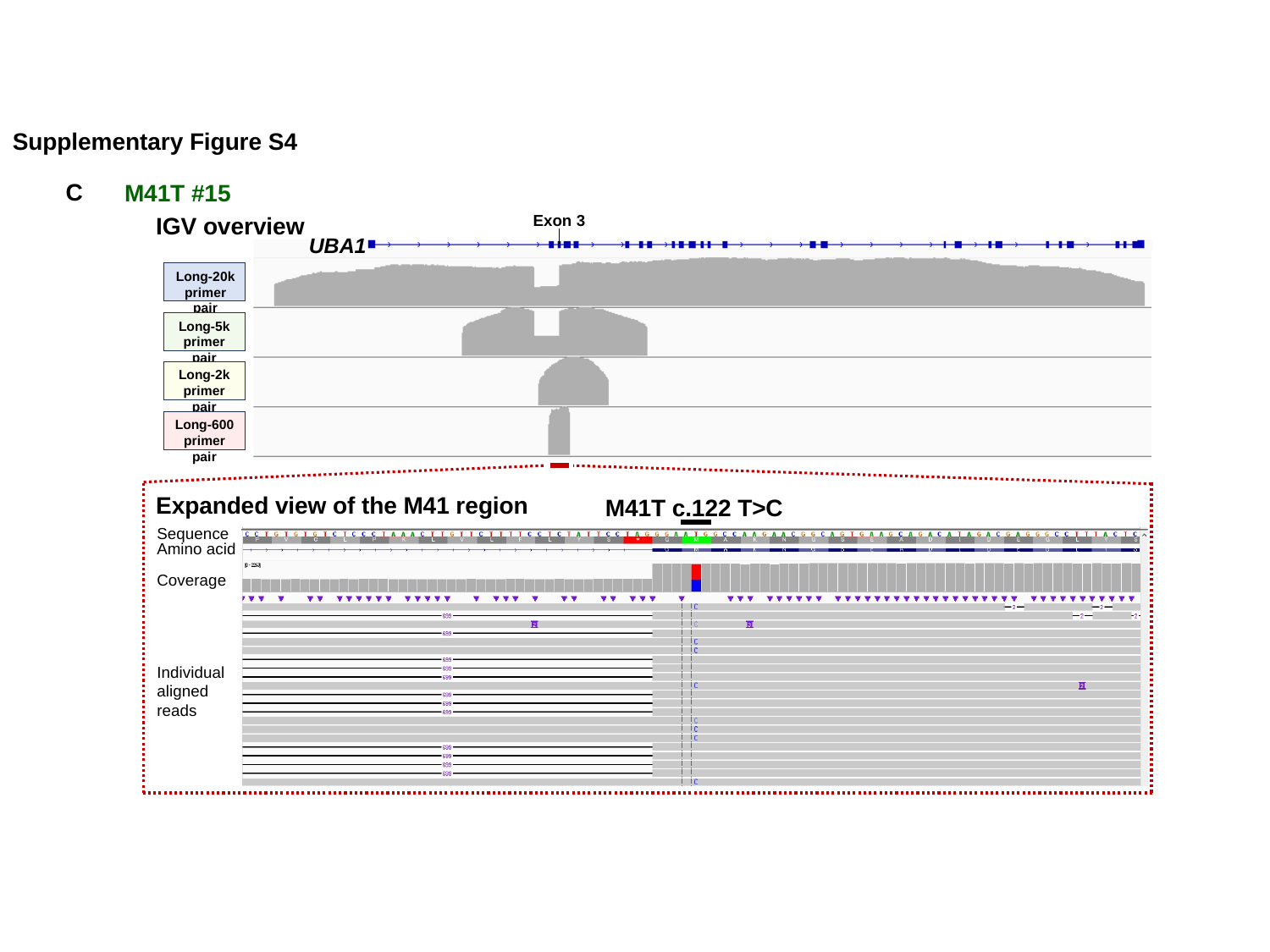

Supplementary Figure S4
C
M41T #15
Exon 3
IGV overview
UBA1
Long-20k
primer pair
Long-5k
primer pair
Long-2k
primer pair
Long-600
primer pair
Expanded view of the M41 region
M41T c.122 T>C
Sequence
Amino acid
Coverage
Individual
aligned reads

### Slide 7
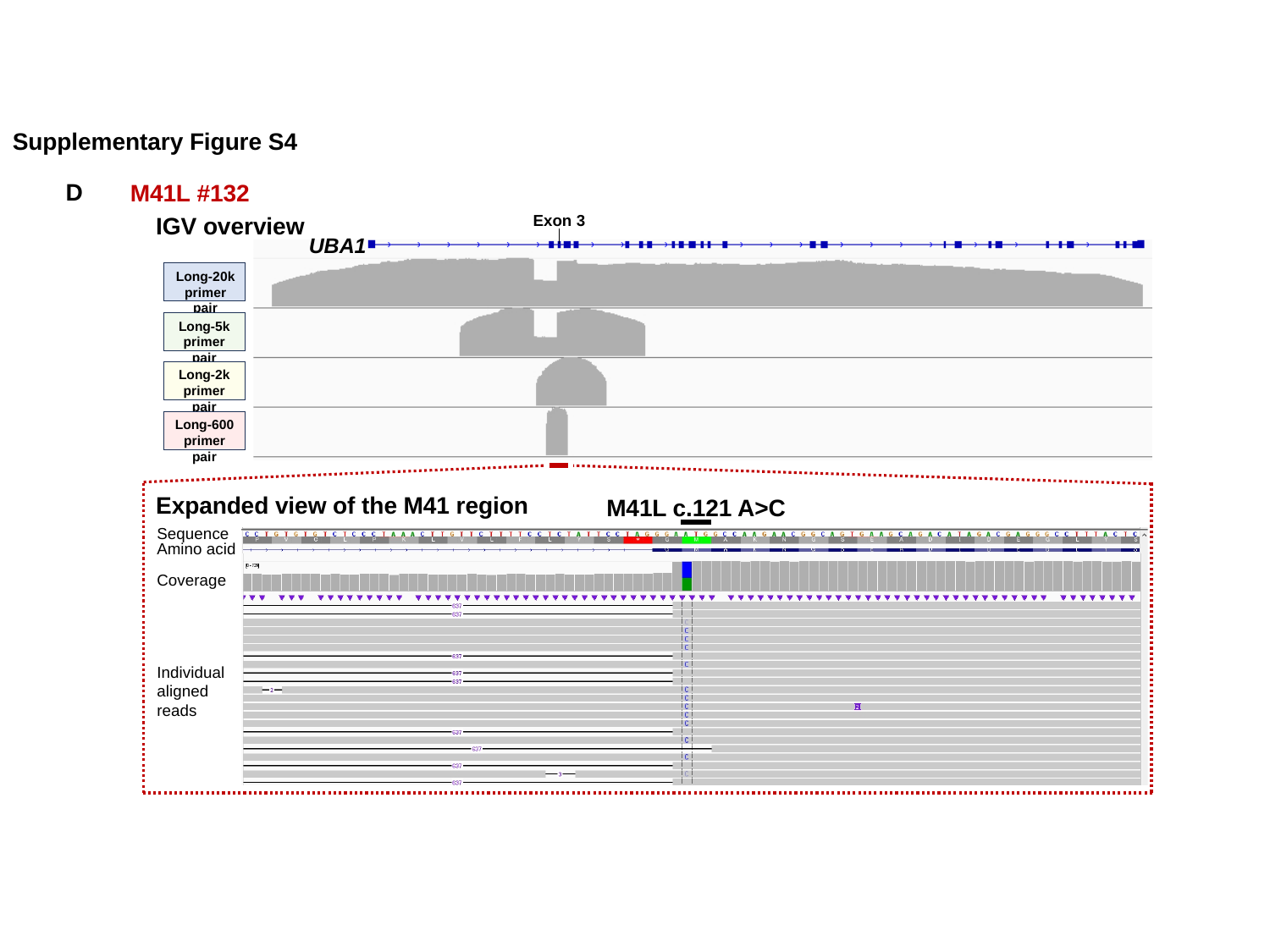

Supplementary Figure S4
D
M41L #132
Exon 3
IGV overview
UBA1
Long-20k
primer pair
Long-5k
primer pair
Long-2k
primer pair
Long-600
primer pair
Expanded view of the M41 region
M41L c.121 A>C
Sequence
Amino acid
Coverage
Individual
aligned reads
